# In vivo spatial coordination with synthetic paracrine signaling

**DOI:** 10.64898/2026.06.26.734902

**Authors:** Kaiwen Luo, Yitong Ma, Hongyi R. Li, Hengyu Li, Mengziang Lei, Margaret B. Swift, Nathan F. Dalleska, Abdullah S. Farooq, Ernesto Criado-Hidalgo, Ann Liu, Mikhail G. Shapiro, Michael B. Elowitz

## Abstract

The immune system uses paracrine signaling to spatially confine potent responses such as inflammation. A bio-orthogonal synthetic paracrine system could enable engineering of analogous multicellular circuits in which different cell types coordinate their functions in a spatially organized fashion. Here, using the plant hormone auxin as a bio-orthogonal chemical signal, we introduce programmable paracrine circuits that distribute sensing and effector functions to different cell types to spatially restrict responses in mouse xenografts. Cells engineered to express auxin biosynthetic genes generated auxin-dense regions with tunable length scales in vivo. This localized signaling ability enabled design of a multicellular sentinel-effector system, in which THP-1 sentinel cells conditionally produce auxin in regions expressing the tumor-specific antigen EGFRvIII, and Jurkat effector cells respond by locally modulating the activity of a chimeric antigen receptor (CAR). This two-cell type system was able to achieve localized activation of engineered effector cells in vivo. These results establish a foundation for engineering multicellular therapeutic systems that focus responses in specific tissue contexts or disease sites.

## INTRODUCTION

The immune system concentrates its most potent inflammatory activities at the specific locations where they are needed. This spatial specificity is essential to minimize tissue-damaging inflammation in healthy bystander regions^1^. Spatial specificity is achieved in part through paracrine communication: sentinel cells such as macrophages detect threats and release diffusible cytokines (e.g., TNFα) that form local gradients to recruit and activate effector cells, such as neutrophils, at diseased sites^2,3^. By contrast, engineered cell therapies generally lack intrinsic mechanisms to spatially confine their activity once activated. In CAR-T therapies, this can cause on-target, off-tumor toxicity when the target antigen is expressed in healthy tissues. For example, HER2 is overexpressed in some tumors but also present in certain normal tissues away from the tumor site, resulting in severe toxicities in clinical trials of HER2-targeting CAR-T cells^4^. The inability to program spatial awareness in cells also limits the use of disease-specific but heterogeneous antigens as targets. For example, the tumor-specific antigen EGFRvIII is absent from normal tissues, providing strong tumor specificity, but it is expressed in only a fraction of tumor cells^5^, preventing its use as a primary target. The ability to spatially restrict CAR-T activity to the tumor region using EGFRvIII could relax the specificity requirements for the CAR. Specifically, it could enable targeting of lower-specificity antigens, such as HER2, within the tumor neighborhood. To enable programming of these and other spatially organized behaviors in vivo, a synthetic, bio-orthogonal paracrine signaling system is needed.

The ideal synthetic paracrine system would function independently of host pathways, signal diffusively through living tissues over tunable length scales, and provide sufficient sensitivity to gate cellular activities within a limited range of signal-sending cells. However, achieving these properties in vivo is challenging. Dense extracellular matrix, complex vasculature and cell movement can all distort signal distribution and strength, while overly strong signaling may reduce spatial confinement. Furthermore, the ability for one cell type to spatially restrict an activity in a confined region requires that a signal propagate from one cell to another and effectively regulate effector activities in the other cell. Inefficiencies at any step in this process could limit overall performance and compromise spatial confinement.

Previous efforts have taken initial steps towards a synthetic, bio-orthogonal paracrine signaling system that can operate in vivo. Synthetic receptor platforms such as GEMS, LIDAR and MESA enable mammalian cells to respond to defined ligands. However, they did not constitute a sender-receiver paracrine system based on bio-orthogonal signals secreted by engineered cells^6–8^. Other studies achieved engineered diffusible communication in vitro, but did not establish spatially restricted function in vivo^9,10^. In contrast to these cell culture studies, other researchers repurposed acetaldehyde and L-tryptophan as synthetic signals in mice. However, these molecules lacked bio-orthogonality and signal localization^11,12^. As a result, a synthetic paracrine system capable of spatially restricting activities among distinct engineered mammalian cell types in vivo remains lacking.

The plant hormone auxin (indole-3-acetic acid, IAA) could provide a useful bio-orthogonal communication channel in engineered mammalian cells^13^. Auxin is well-tolerated in human studies (e.g., ~100 mg/kg with no reported acute adverse effects)^14,15^. Further, mammalian cells can be engineered to produce auxin from tryptophan by expressing a two-gene biosynthetic pathway imported from bacteria. Secreted auxin can then be sensed using the osTIR1 F-box protein, which causes degradation of proteins tagged with the cognate auxin-inducible degron (AID)^13,16,17^. This sender-receiver system generated long-range (centimeter scale) spatial gradients of auxin in cell culture^13^. However, it has remained unclear whether it could function over relevant length scales in vivo. In addition, it is also essential to engineer useful responses to signal detection. This requires linking sensing of an orthogonal signal to control of a functional component, such as a CAR. Yet the ability to generate, sense, and respond to local bio-orthogonal signals in vivo has not been demonstrated.

Here, we introduce a synthetic paracrine signaling system that confines cell activities to limited spatial regions in living tissues (**Fig. 1**). We first engineered auxin-producing cells that generate localized, dose-dependent paracrine signaling between CHO-K1 sender and receiver cells in mouse xenografts over scales of hundreds of microns to several millimeters. Using this signaling system, we designed an immune-inspired two-cell-type synthetic paracrine (“synthacrine”) circuit that links antigen sensing to an effector response (**Fig. 1A**). In this circuit, sentinel-like cells secrete auxin in response to a tumor-specific antigen, while engineered Jurkat cells permit CAR activity in response to auxin signaling (**Fig. 1C**). Remarkably, the two cell type system successfully restricted Jurkat activation to tumor regions in vivo. More generally, these results demonstrate how a bio-orthogonal paracrine communication channel can function predictably in vivo, providing a foundation for future spatially aware multicellular circuits that help focus therapeutic functions in complex tissue environments. This capability may also enable engineering of other multicellular, immune-like systems.

**Figure 1:**
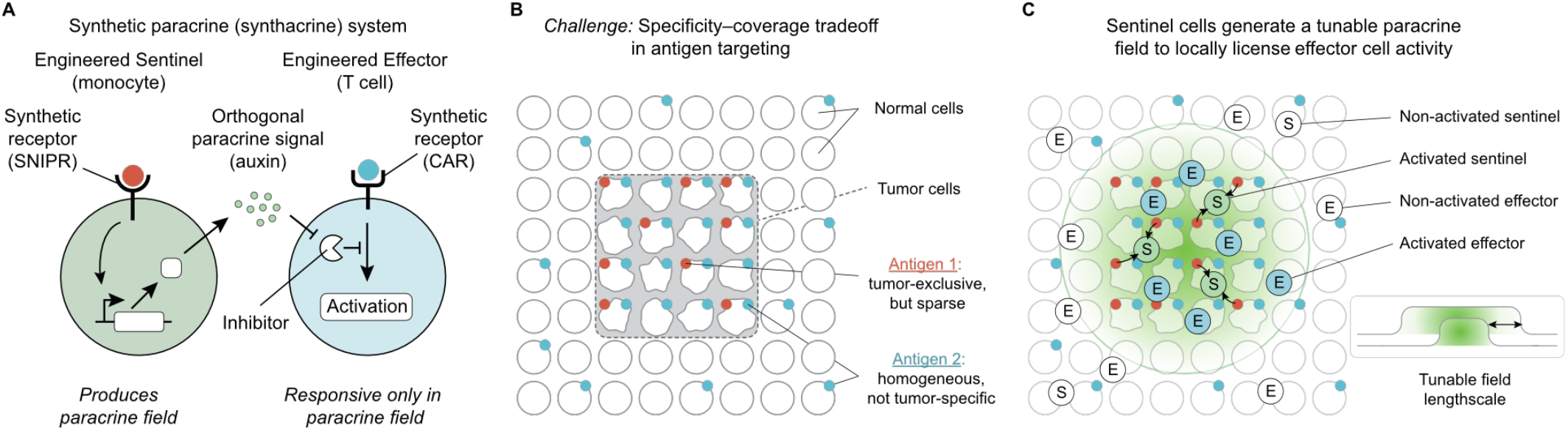
A synthetic paracrine signaling system could restrict effector activity to defined spatial regions. (**A**) Design of a synthetic paracrine system. In this system, engineered sentinel cells (green, left) respond to an antigen (red circle) by producing a diffusible small-molecule signal (green dots). The signal de-inhibits effector cells (blue, right) within a local area. Examples of the specific components used here are indicated in parentheses. (**B**) A cancer-inspired design challenge: tumor antigen targeting can face a specificity-coverage tradeoff, when one antigen (antigen 1, red dots) is tumor-specific but spatially heterogeneous, while a second (antigen 2, blue dots) is homogeneous within the tumor but not tumor-specific. Targeting either antigen alone would lead to incomplete response or on-target, off-tumor toxicity, respectively. (**C**) The synthetic paracrine system addresses this challenge. Sentinel cells recognize antigen 1, and locally secrete the bio-orthogonal paracrine signal to “paint” the tumor region. This paracrine signaling field, in turn, locally licenses effector cells targeting antigen 2. Critically, the size of this field can be tuned to match the desired spatial range.

## RESULTS

### A two-cell synthetic paracrine system achieves local, dose-dependent signaling in living tissues

As a first step towards establishing a synthetic paracrine system that can operate in living tissues, we engineered cells that send or receive auxin signals. To create a constitutive auxin sender cell, we stably integrated the auxin biosynthetic genes iaaM and iaaH in CHO-K1 cells. To create auxin receiver cells, we co-expressed the auxin receptor F-box protein osTIR1, an mCherry-AID reporter (auxin-degradable mCherry), and a constitutive BFP lineage marker in CHO-K1 cells (**Fig. 2A**). The resulting cell line exhibited a monotonic ~30-fold decrease in mCherry fluorescence over an auxin titration (**Fig. S1A, right**).

**Figure 2:**
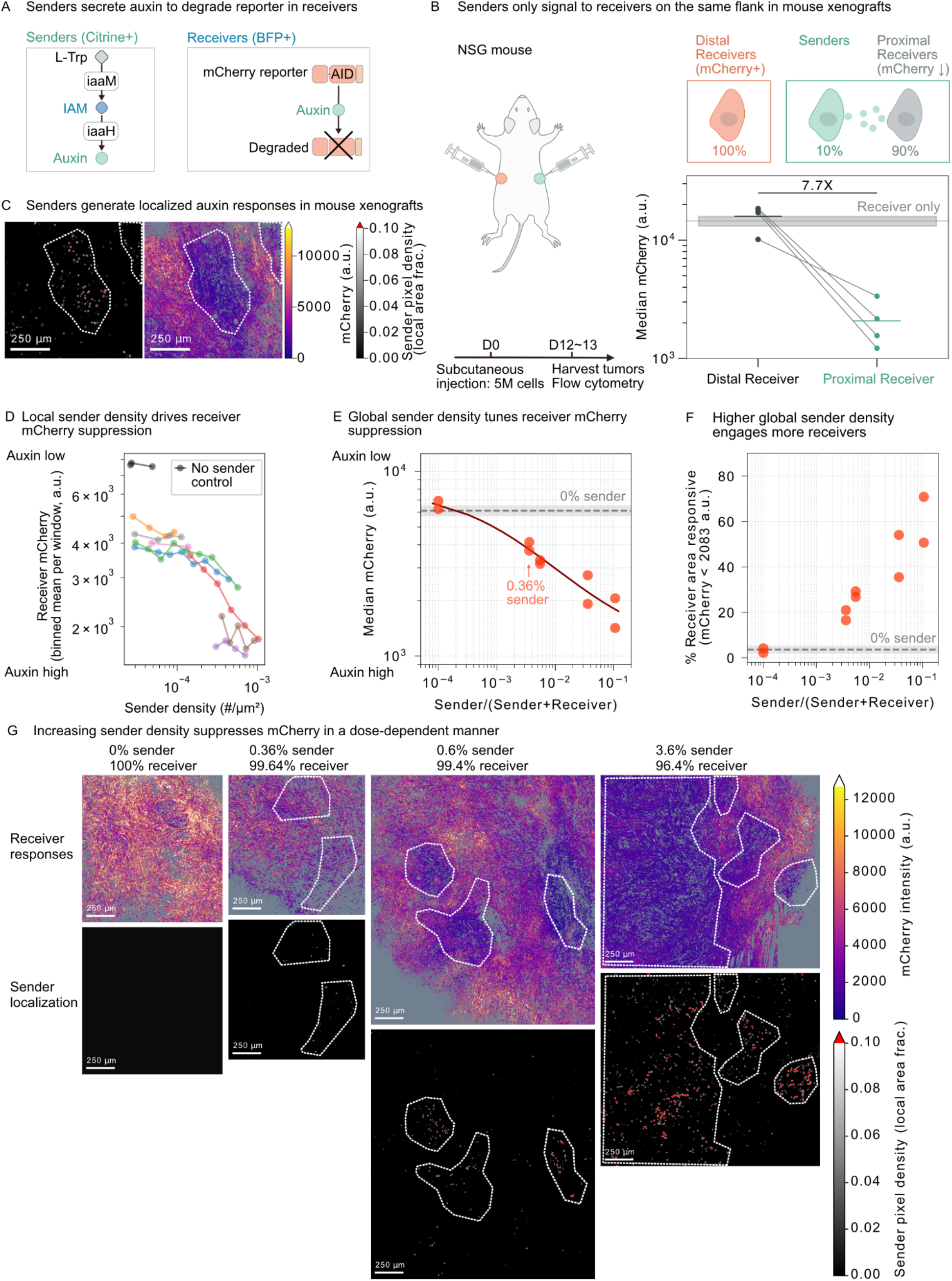
A two-cell synthetic paracrine system achieves local, dose-dependent signaling in living tissue. (**A**) Auxin “sender” cells synthesize auxin from L-tryptophan (L-Trp) via a two-step pathway. Auxin diffuses to “receiver” cells, where it promotes osTIR1-dependent degradation of a mCherry-AID reporter. AID, auxin-inducible degron. (**B**) Bilateral xenograft assay to test spatially confined signaling. One flank contained a receiver-only xenograft; the contralateral flank contained a mixed xenograft (10% senders, 90% receivers). Receiver median mCherry fluorescence was used as a proxy for auxin exposure (lower mCherry indicates higher auxin). Gray dotted line and shaded band indicate receiver mCherry in a control mouse bearing a single receiver-only xenograft (mean ± s.d.). N = 4 mice. Horizontal lines indicate mean values. (**C**) Confocal tile-scan mosaics of CHO-K1 xenograft sections were processed to generate spatially matched maps of receiver response and local sender abundance (representative cropped regions shown). Processed images are for display only and were not used for quantitative analysis. Receiver pixels were identified from BFP signal and displayed as a heat map of mCherry intensity (over-range values indicated as white). Sender pixels were defined as GFP+/BFP-/mCherry-. The sender pixel map was Gaussian-smoothed to yield a local sender abundance map (over-range values indicated as red). Scale bars, 250 μm. Dotted lines indicate sender-rich regions for visual comparison. Because these maps were generated from single-plane confocal images, sender cells located outside the imaging plane are not represented. (**D**) Local sender density–receiver response relationship from images of xenograft sections, computed in 200 µm × 200 µm windows. For each window, sender density and median receiver mCherry were calculated; windows with ≥ 50 receivers were retained. Window-level values were log-binned by sender density and plotted with one curve per sample instance. Most curves represent independent xenograft samples (N = 7 independent samples total); one sample is shown as two curves from anatomically distinct xenograft regions. No-sender controls (receiver-only, 0% sender, and 0.01% sender seeded with sender-free fields by visual inspection) are shown in black to denote baseline mCherry; rare detected “senders” likely reflect classifier noise near the detection limit. (**E**) Receiver median mCherry across xenografts seeded with varying sender fractions (receiver definition as in panel **C**). “0% sender” is a receiver-only control xenograft; horizontal line and shading indicate median ± median absolute deviation (MAD). A Hill function was used to fit data points (**Methods**). See **Supplementary Data** for fitted parameters. N = 2 xenografts for each sender-fraction group. (**F**) Fraction of signal-responsive receiver area across xenografts seeded with varying sender fractions (quantified using pixels). Receiver pixels were defined using the same classification as panels C and E. Responsive receiver pixels were defined using a response threshold set from receiver-only controls and then applied uniformly across samples. Horizontal line and shading indicate receiver-only control median ± MAD. N = 2 xenografts for each sender-fraction group. (**G**) Representative xenograft section images (confocal; processed) for xenografts seeded with different sender fractions (processed as in panel **C**). Dotted lines indicate sender-rich areas for visual comparison.

To test whether these cells could productively signal in vivo, we coinjected the CHO-K1 senders and receivers subcutaneously into immunocompromised NOD *scid IL-2Rγ*^−*/*−^ (NSG) mice to form mixed xenografts (**Fig. 2A and 2B**). We allowed the xenografts to develop for 12-13 days, sacrificed mice, and imaged sender and receiver cells by confocal fluorescence microscopy of xenograft slices and performed flow cytometry of live cells extracted from the xenografts (**Methods**).

Auxin signaling was local, rather than systemic. In a bilateral xenograft model, receivers on one flank were unaffected by senders on the opposite flank (**Fig. 2B**). Further, liquid chromatography-mass spectrometry (LC-MS) analysis revealed that serum auxin levels from mice with the sender/receiver xenograft were not elevated compared to background levels, and established an upper limit of ~0.02 µM for systemic auxin concentration generated by sender xenografts under these conditions (**Fig. S1A, Methods**). These results are consistent with reports of a short (~20-70 min) in vivo half-life of auxin after a single injection in mice (250 mg/kg)^18^, and renal/hepatic clearance of auxin^19^. Together, these results suggest that auxin can provide transient, spatially restricted signaling in vivo over organismal length scales.

To understand how local auxin signaling is in vivo, we systematically varied the initial fraction of sender cells within mCherry reporter xenografts and analyzed them by confocal imaging. We analyzed mCherry fluorescence in regions containing elevated densities of sender cells. These regions exhibited reduced mCherry fluorescence, consistent with in vivo paracrine signaling (**Fig. 2C**). To quantify the relationship between local sender density and reporter fluorescence, we partitioned confocal images into 200 µm × 200 µm windows. For each window containing at least 50 receiver cells, we computed the 2-D sender density and median receiver mCherry. We then plotted the median receiver mCherry in each window versus the sender density in the same window, averaged in bins of similar sender density, for multiple samples. As controls, we analyzed samples lacking sender cells altogether. We observed progressively stronger suppression with increasing local sender density (**Fig. 2D**). This reduction extended to low sender densities of ~40 cells/mm^2^, or 4×10^−5^ cells/µm^2^. By contrast, receiver-only xenografts (0% senders) and low-sender xenografts (0.01% sender seeded; sender-free by visual inspection in the analyzed fields) exhibited no reduction in mCherry (**Fig. 2D**). Together, these data suggest that sender cells can locally suppress receiver mCherry fluorescence in a density-dependent fashion.

Strong suppression of receiver mCherry fluorescence throughout the tumor could be achieved with modest sender cell fractions. To quantify receiver mCherry fluorescence in images of xenograft sections, we first classified receiver pixels (BFP+) and then computed the median mCherry intensity across all receiver pixels within each xenograft. This median receiver-pixel mCherry intensity was compared across xenografts seeded with different sender fractions. Receiver-only (0% sender) xenografts provided a baseline. Increasing sender cell fraction led to a monotonic dose-dependent reduction in global receiver mCherry intensity (**Fig. 2E**). Sender fractions as low as ~0.36% produced measurable reductions in receiver mCherry compared to receiver-only controls (**Fig. 2E**). To examine this effect spatially, we measured the fraction of receiver area that responded at each sender fraction. We classified BFP+ receiver pixels as responsive if the mCherry intensity of the same pixel fell below an auxin-response threshold (**Methods**). This analysis method had a low background, exhibiting negligible responses in negative (0% sender) controls (**Methods**). For each xenograft section, we also quantified the responding area fraction, defined as the proportion of receiver pixels below the auxin-response threshold (**Methods**). This responding fraction increased steadily with increased sender fraction (**Fig. 2F, 2G**). At the highest sender percentage examined (3.6–10%), responding regions extended to millimeter-scale domains. For example, a global sender fraction of 10% affected up to ~60% of the receiver area (**Fig. 2F, 2G**). Thus, relatively small sender fractions can effectively signal across larger tumor regions.

Together, these results establish auxin as an orthogonal paracrine signal that supports local, dose-dependent communication in living tissue. Within the sender/receiver xenograft, clusters of senders generate “response domains” whose spatial extent scales with sender density, ranging from small clusters to contiguous regions spanning hundreds of micrometers to millimeters (**Fig. 2C, 2F, 2G**). Thus, modest sender fractions (3.6–10%) can drive broad receiver suppression, while maintaining local containment and minimal systemic spillover.

### Engineered sentinel cells conditionally secrete auxin in response to tumor-specific antigens

We next asked whether engineered “sentinel” cells could detect heterogeneous tumor-specific antigens like EGFRvIII and label the surrounding region with auxin to spatially gate the activity of T cells (“effectors”) (**Fig. 1 and Fig. 3A**). This requires circuits that conditionally activate expression of auxin biosynthetic enzymes in response to sensing of EGFRvIII. To implement this, we designed and optimized an antigen-gated THP-1 sender cell (“sentinel”) that uses an EGFRvIII-responsive SNIPR to activate expression of auxin biosynthetic enzymes.

**Figure 3:**
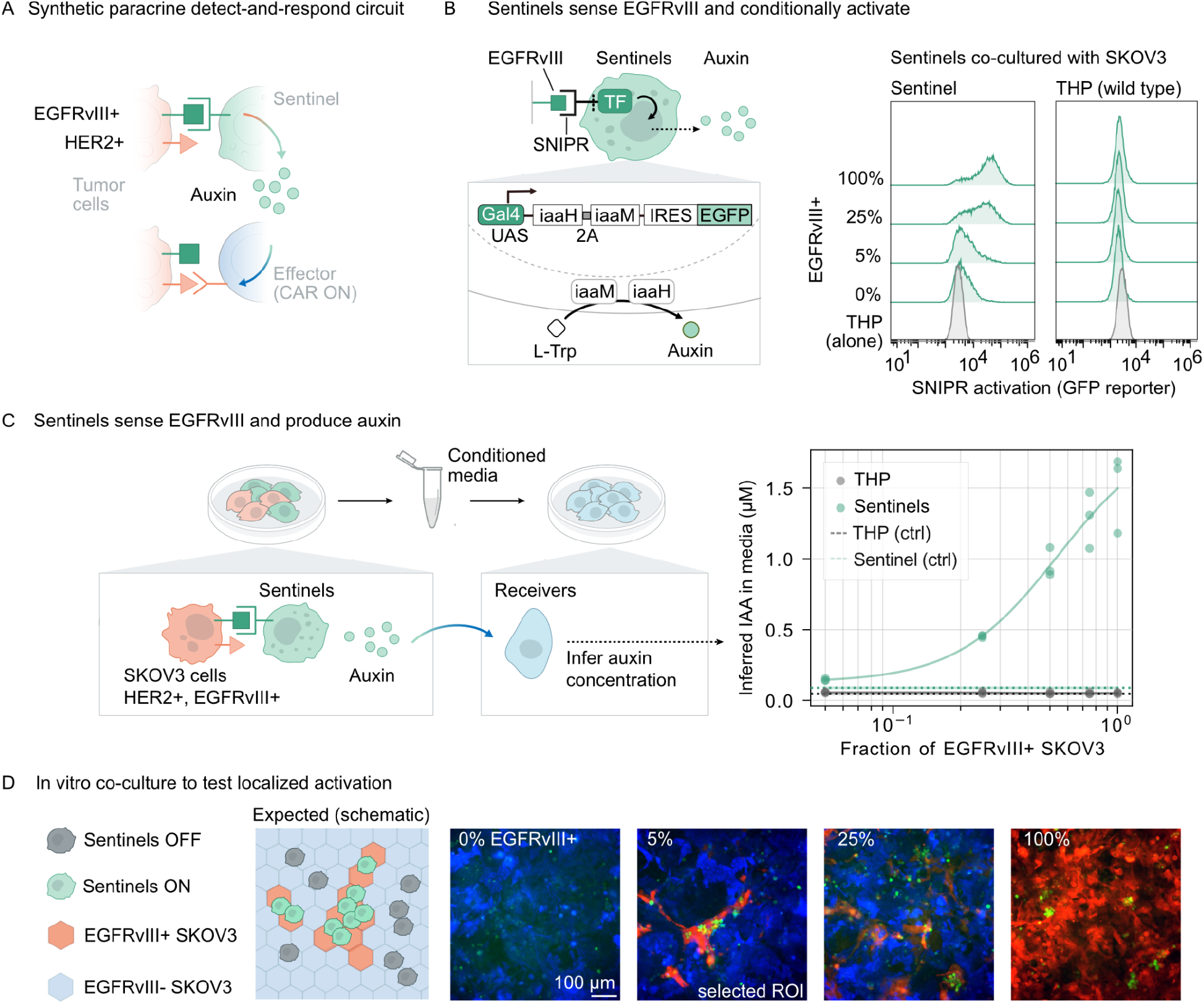
Engineered sentinels secrete auxin in response to EGFRvIII. (**A**) Synthetic paracrine circuit design. THP-1 monocyte “sentinel” cells detect EGFRvIII and secrete auxin. Jurkat “effector” cells express an auxin-gated anti-HER2 CAR, and respond to HER2+ targets only when auxin is present. (**B**) EGFRvIII recognition by SNIPR triggers release of a synthetic transcription factor that induces iaaM/iaaH expression (auxin biosynthesis). Sentinels were co-cultured with SKOV3 cells containing varying fractions of EGFRvIII+ cells and analyzed by flow cytometry; activation was reported by a co-expressed EGFP marker. (**C**) Auxin output as a function of EGFRvIII+ fraction. Conditioned media from sentinel-SKOV3 co-cultures were applied to auxin receiver cells to infer auxin concentration, using a side-by-side IAA standard curve. “Sentinel (ctrl)” and “THP (ctrl)” denote co-culture with EGFRvIII-(wild-type) SKOV3, where horizontal lines and shading indicate median ± MAD. A Hill function was used to fit data points (**Methods**). See **Supplementary Data** for fitted parameters. N = 3 for each condition. (**D**) Representative co-culture images. Activated sentinels (EGFP+, green) preferentially co-localize with EGFRvIII+ SKOV3 (mScarlet+, red) rather than EGFRvIII-SKOV3 (BFP+, blue) across input mixtures. IAA, indole-3-acetic acid (auxin).

We selected the monocyte cell line THP-1 as the chassis for the sentinel cells, motivated by the natural role of macrophage-lineage cells in surveying tissue microenvironments and integrating local inflammatory signals, as well as their ability to infiltrate and persist in solid tumors^20–22^.

We first optimized the auxin biosynthetic constructs to boost their expression and activity in THP-1 cells. We considered a variety of expression construct configurations for the auxin biosynthetic genes. One configuration generated an elevated auxin production rate (constitutive THP senders; **Fig. S2A, Methods**). Supplementation of auxin precursor indole-3-acetamide (IAM) eliminated the differences in auxin production rates among the designs, suggesting that iaaM, which catalyzes conversion of Trp to IAM, is rate-limiting for auxin production (**Fig. S2B**). Therefore, the gain in auxin production in this configuration likely resulted from higher iaaM expression, possibly because of its earlier position in the P2A peptide series^23^.

To engineer sentinel cells capable of sensing a tumor-specific antigen, we used lentiviral constructs to integrate an EGFRvIII-responsive SNIPR (**Methods**)^9^ that releases the Gal4 transcription factor when the receptor engages with EGFRvIII (**Fig. 3B**). We also incorporated a Gal4-activated construct co-expressing the auxin biosynthetic enzymes iaaH and iaaM together with EGFP, thus creating THP sentinel cells that secrete auxin only after detecting EGFRvIII in the microenvironment.

Next, we tested the ability of the engineered sentinel THP-1 cells to respond to EGFRvIII+ SKOV3 human ovarian cancer cells in a co-culture. We varied the fraction of EGFRvIII+ SKOV3 target cells while holding the total SKOV3 cell number constant, and quantified EGFP expression in the co-cultured sentinel cells. EGFP signal increased with antigen input, whereas wild-type THP-1 lacking SNIPR remained unchanged across conditions (**Fig. 3B**). Functionally, auxin in supernatant increased with the EGFRvIII+ fraction, reaching ~1.5 µM when 100% of SKOV3 target cells were EGFRvIII+ (**Fig. 3C**). This result indicates that antigen engagement can drive functionally relevant levels of auxin production over the time scale of 3-4 days (**Methods**).

To operate as a paracrine system, the sentinel cells should be activated only in proximity to EGFRvIII+ tumor cells. To assess this spatial restriction of sentinel cell activation, we co-cultured sentinel cells with a monolayer of SKOV3 cells. The monolayer contained patches of EGFRvIII+ cells interspersed with EGFRvIII-cells (**Fig. 3D**). We imaged the co-culture 5 days later by fluorescence microscopy. Activated sentinel cells co-localized with EGFRvIII+ (mScarlet+) tumor cells rather than EGFRvIII-(BFP+) neighbors (**Fig. 3D**), likely due to decreased mobility of sentinels upon antigen detection. Taken together, these data indicate that engineered THP-1 sentinel cells can activate locally near EGFRvIII+ cells and secrete auxin in a dose-dependent manner in cell culture.

### Auxin gates CAR function in engineered effector cells

Localized auxin production opens up the possibility of engineering receiver cells with spatially restricted activities. Engineered T cells represent a major platform for cell therapy. Restricting their activities to disease sites could mitigate issues such as on-target, off-tumor toxicity when targeting tumor-associated antigens with lower specificity, such as HER2^4^. As a demonstration of spatially restricted activity, we sought to engineer auxin-dependent CAR signaling in Jurkat T cells (**Fig. 4A**), and used CD69 induction as a tractable readout of effector activation rather than tumor killing.

**Figure 4:**
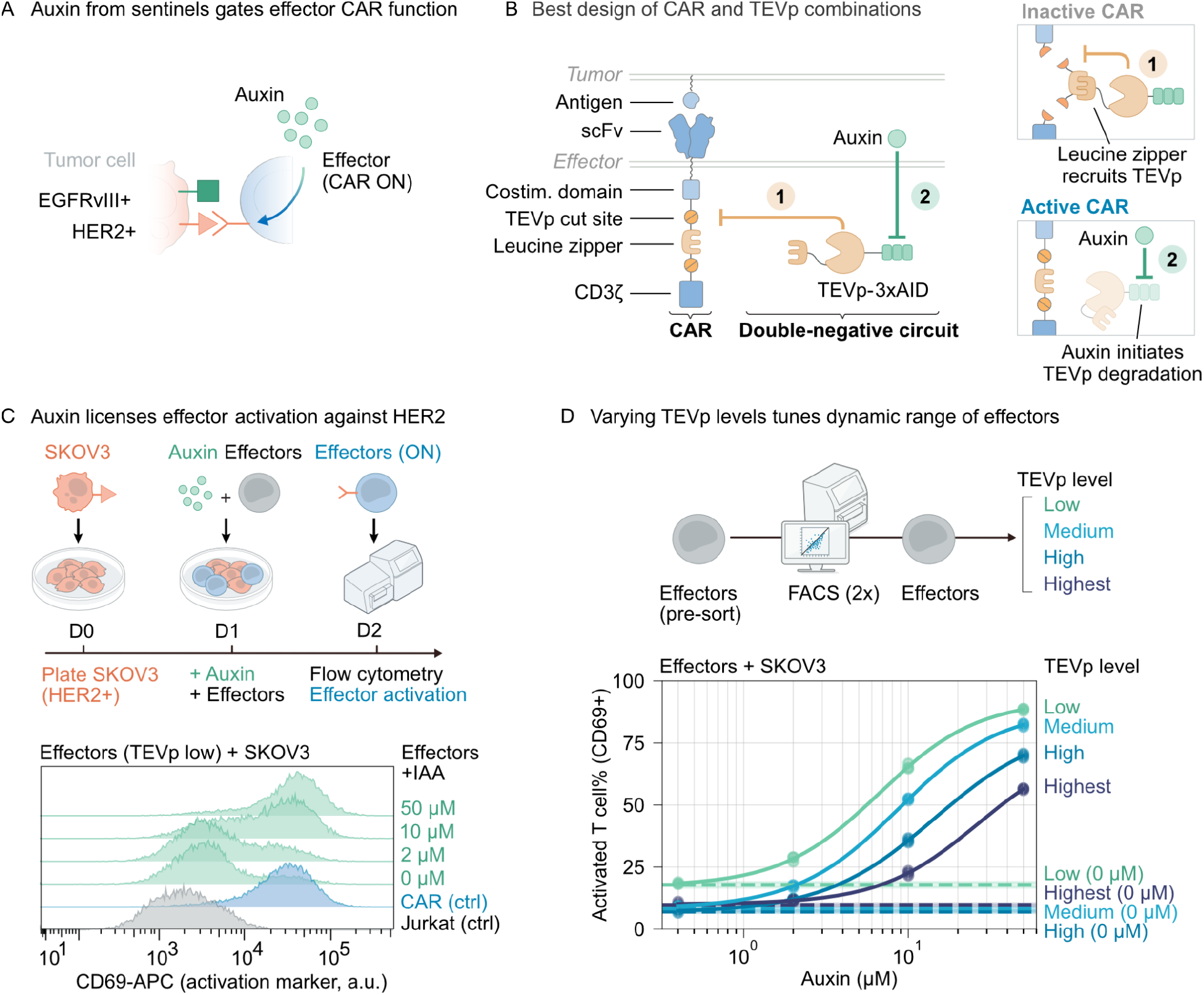
Engineered effectors couple auxin to CAR activity. (**A**) Effector module in the three-cell circuit: Jurkat cells sense auxin to permit functional anti-HER2 CAR activity against target cells. (**B**) Auxin-gated CAR design. TEV protease (TEVp) suppresses CAR activity, while auxin promotes CAR function by degrading TEVp. TEVp is fused to a tandem auxin-inducible degron (3xAID) at the N-terminus, and a C-terminal leucine zipper to promote interaction with the CAR. The CAR contains a matching leucine zipper flanked by two TEVp cleavage sites, enabling TEVp-dependent inactivation. (**C**) Auxin permits effector activation. Effector cells were co-cultured with tumor cells (1:2 ratio) for 24 h with indicated IAA doses, then analyzed with flow cytometry; CD69 was used as the activation readout. Effectors were sorted to have low TEVp and high CAR expression (see panel **D**). (**D**) Tuning auxin sensitivity by TEVp expression. Effectors were sorted into bins by TEVp/EGFP level (see **Fig. S3B**) and assayed as in panel **C** to quantify auxin-dependent activation across TEVp expression regimes. Activation fractions of effector cells from no-auxin control groups were labeled “(0 µM)”, where horizontal lines and shading indicate mean ± s.d.. A Hill function was used to fit data points (**Methods**). See **Supplementary Data** for fitted parameters. N = 3 for each condition. IAA, indole-3-acetic acid (auxin).

Engineered auxin-dependent regulation typically involves incorporation of auxin-inducible degrons (AIDs) on target proteins, leading to negative regulation of the target protein by auxin. To invert the sign of regulation, and thereby make auxin a permissive signal, we designed a double-negative synthetic protein-level circuit (**Fig. 4B**)^24,25^. In this design, auxin destabilizes a constitutively expressed viral TEV protease (TEVp), which otherwise cleaves engineered TEV sites within the CAR to inactivate it. With this design, auxin produced by sentinel cells should suppress TEVp and thereby permit CAR function in localized regions (**Fig. 4A**).

To engineer this system, we systematically explored a variety of CAR and TEVp variant designs. Each CAR design incorporated TEVp cleavage sites or TEVp degrons at various numbers and locations within the protein (**Fig. S3C-G**). In some cases, we split the CAR into two distinct chains with the goal of increasing regulability (**Fig. S3E**). Similarly, each TEVp variant incorporated different numbers of AIDs at different sites within the protein, or was split into two halves to improve dynamic range (**Fig. S3D-G**). Finally, we analyzed system designs in which both the CAR and TEVp were tagged with complementary leucine zippers to facilitate interactions (**Fig. 4B and S3D-G**). For each system design, we quantified three operating states: basal CAR function in the absence of TEVp, TEVp-mediated suppression, and auxin-dependent rescue via TEVp degradation. We assayed output by co-culturing engineered Jurkat cells with HER2+ SKOV3 targets, and measuring CD69 induction after 24 h, using 50 µM IAA to approximate maximal de-repression (**Methods, Fig. 4C**).

Many designs exhibited low CAR activity, weak or no TEVp dependence, or minimal response to auxin (**Fig. S3C-F**). However, some designs worked well (**Fig. S3G**). The best-performing configuration included leucine zippers for CAR-TEVp interaction, a tandem fusion of 3 AIDs at the N-terminus of TEVp, and, within the CAR, two TEVp cleavage sites flanking the leucine zipper between the 4-1BB and CD3ζ domains (**Fig. 4B and S3G**). We further sorted the Jurkat cells engineered with the optimal design for different CAR/TEVp expression regimes and identified a regime that yielded good sensitivity to auxin (sorted cells denoted as “effectors”, here with low TEVp expression). From 0–50 µM IAA, the percentage of CD69+ effectors increased dose-dependently and, at 50 µM, reached a level comparable to the CAR-only positive control (**Fig. 4C**). We note that co-expression of EGFRvIII in SKOV3 did not alter auxin-dependent activation at matched IAA doses and cell ratios, allowing direct comparison of circuit behaviors when co-cultured with SKOV3 cells with or without EGFRvIII (**Fig. S3A**).

We next screened cells expressing components at varying levels to identify high-performing cells. Using a co-expressed EGFP reporter, we sorted effector cells with different levels of TEVp, and analyzed their response to auxin titration (**Fig. 4D and S3B**). Low-TEVp-expressing cells attained high CD69 induction at lower (~10 µM) auxin concentrations, with only modest background signaling. By contrast, intermediate TEVp levels minimized background activity and improved the dynamic range (**Fig. 4D**). The highest levels of TEVp expression required higher auxin concentrations to respond, while maintaining similar levels of background activity suppression. These results are consistent with TEVp abundance shifting the effective thresholds for auxin sensing (**Fig. 4D**).

Taken together, these results indicate that auxin-gated CAR effectors are sensitive and tunable. Modulation of TEVp expression enables matching of operating points to sentinel-derived auxin levels. This module should therefore support spatial restriction of effector activity when paired with auxin-secreting sentinels.

### The sentinel-effector system conditionally detects tumor-specific antigen in vitro

To achieve spatial gating of CAR activity, signals must propagate through three sequential steps: sentinels must sense EGFRvIII expressed on target cells, produce auxin, and induce auxin-dependent de-inhibition of CARs in effectors. Inefficiencies at any step in this process, including insufficient sensing or production of auxin, or incomplete de-inhibition of the CAR, limit the overall responsiveness of the sentinel-effector system.

We therefore set out to test whether the circuit had sufficiently low background activation and sufficiently high on-target activation to function effectively. We established a three-component co-culture, comprising SKOV3 target cells, THP-1 sentinel cells, and Jurkat effector cells. SKOV3 served as both the EGFRvIII source for sentinel activation and the HER2+ target for effector CAR engagement. Low-TEVp Jurkat effector cells were used for better sensitivity at lower auxin concentrations. We plated SKOV3 cells with THP-1 sentinels on day 0 at equal densities. We then added Jurkat effector cells on day 1, at a lower density due to their higher proliferation rate. We also supplemented the media with the auxin precursor IAM (50 µM), as an experimental tuning input to increase auxin output from sentinels (**Fig. S2B**) and probe circuit operating range. After 3 days of co-culture, we quantified T cell activation using CD69 as an activation marker.

Under these conditions, signals successfully propagated through the entire circuit (**Fig. 5**). With the full system, including the EGFRvIII+ SKOV3 cells in the presence of IAM, the fraction of activated effector Jurkat cells increased from <20% (baseline) to 60% compared to a negative control condition lacking EGFRvIII expression on the SKOV3 cells. EGFRvIII levels were not limiting for activity, as the full system reached similar activation levels as observed with positive control sentinel cells that constitutively produce auxin.

**Figure 5:**
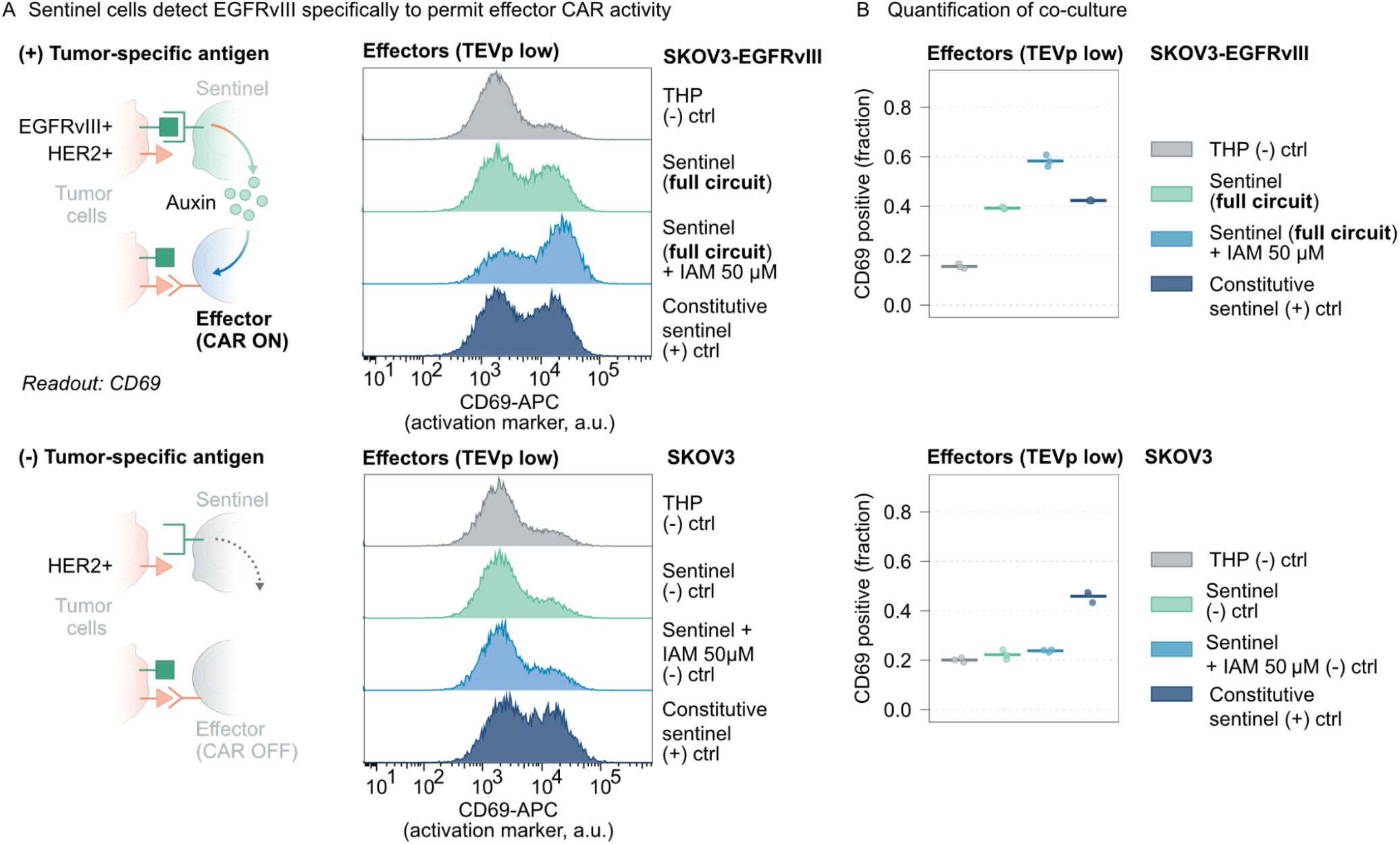
The sentinel-effector circuit conditionally detects tumor antigen in vitro. (**A**) Full-circuit activation measured by co-culture assay (constitutive sentinels are the THP senders, Ver2, from **Fig. S2**). (Top) 25,000 SKOV3-EGFRvIII cells were co-cultured with 25,000 sentinel cells for 1 day, then 6,250 effector cells and IAM were added as indicated. IAM was supplied to boost auxin production (**Fig. S2**). Cultures were maintained for 3 additional days, then analyzed by flow cytometry; CD69 in effectors was used as the readout. (Bottom) Same co-culture setup with EGFRvIII-SKOV3 cells as a negative control. (**B**) Summary across conditions, showing the CD69+ fraction in the effector population from (**A**) plus two additional biological replicates. N = 3 for each condition. Horizontal bars indicate mean values.

In the absence of EGFRvIII, circuit output remained at baseline. With EGFRvIII-SKOV3 targets, Jurkat activation remained ~20% regardless of co-culture with THP sentinels or negative control THP-1 cells, indicating minimal sentinel background, and negligible THP-Jurkat crosstalk (**Fig. 5B, lower;** baseline consistent with **Fig. 4D**).

Auxin production did not interfere with CD69 signaling in wild-type Jurkat or Jurkat with canonical anti-HER2 CAR. As expected, activation of auxin-independent, CAR-expressing Jurkat cells was unaffected by both EGFRvIII and auxin production (**Fig. S4**). Additionally, wild-type Jurkat cells lacking CAR expression were not activated by the circuit under any conditions (**Fig. S4**).

Finally, inclusion of IAM enlarged the dynamic range of the response (**Fig. 5**). In its absence, the fraction of activated Jurkat cells was reduced from 60% to ~40%. Notably, IAM did not increase the background activation (**Fig. 5B, lower**). Thus, inclusion of IAM can enhance circuit responses without additional background.

Taken together, these results show that the multicellular circuit is capable of transmitting information via auxin signaling from sensing of the EGFRvIII tumor antigen by sentinel cells to license anti-HER2 CAR activation in a distinct Jurkat cell population, fulfilling the key requirements for a bio-orthogonal paracrine system.

### The sentinel-effector circuit spatially restricts effector activation in vivo

The three-dimensional in vivo environment differs in numerous ways from two-dimensional cell cultures. For example, the presence of extracellular matrix and vasculature could alter auxin transport, removal, and signaling. Additionally, the migration and proliferation pattern of engineered cells in vivo is not predictable. Therefore, we sought to investigate whether the sentinel-effector circuit can enable localized activation in a murine subcutaneous tumor model.

We subcutaneously co-implanted THP-1 sentinels, Jurkat effectors (Low-TEVp), and SKOV3 targets at a 1:1:1 ratio in NSG mice. In each experiment, we compared one synthetic paracrine circuit with a control condition in the same mouse by implanting different cell combinations on opposite flanks. To increase sensitivity (**Fig. 5**), mice were provided with auxin precursor IAM in PBS for drinking (0.5 mg/mL) on days 4-7. We collected the tumors for analysis on day 7 (**Fig. 6A**). We then cryosectioned tumors, stained for human CD3 (effectors) and human CD69 (activation of effectors), and used fluorescent microscopy to distinguish effectors (CD3+) and EGFRvIII+ tumor cells (mScarlet+) at single-cell resolution. Finally, we quantified effector activation by analyzing CD69 fluorescence images (**Methods**).

**Figure 6:**
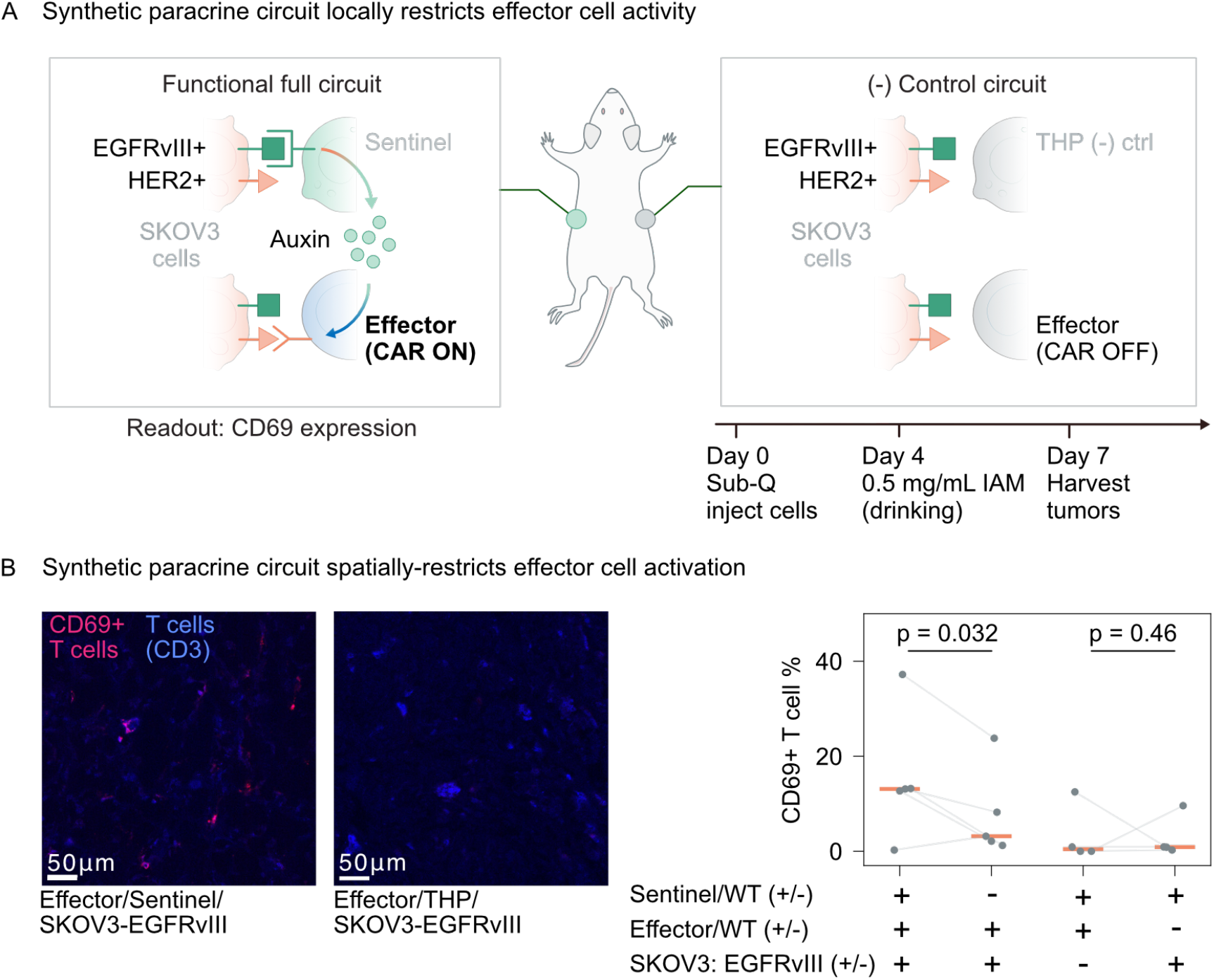
Synthetic paracrine circuit locally restricts engineered T cell activity in vivo. (**A**) SKOV3 cells, THP-1 sentinels, and Jurkat effectors were mixed 1:1:1 and implanted subcutaneously in NSG mice. Full-circuit and negative-control implants were placed on opposite flanks to assess spatial localization in vivo. Effector activation was quantified by CD69. Four days post-implantation, mice were provided IAM in PBS for drinking to boost sentinel auxin production. Seven days after implantation, tumors were harvested, and sectioned for staining and imaging. (**B**) (Left) Representative confocal images of fixed tumor sections stained for effectors (CD3/Pacific Blue, blue) and activation (CD69/APC, red). The processed images are for display only; further quantification was performed on unprocessed data using predefined thresholds described in **Methods**. Linear display ranges were adjusted uniformly across both entire images; the maxima for the blue/CD3 and infrared/CD69 channels were set to 1,000 and 15,823 a.u., respectively, based on a no-primary-antibody control. The images are tile-scan mosaics, acquired automatically and stitched within imaging software. Scale bar, 50 µm. (Right) Quantification of confocal images across tumors/mice for CD69+ T cell %, after segmentation to identify T cells. CD69+ cells were defined as those with mean infrared intensity > 5,000 a.u., a threshold that yields near-zero positives in the negative control (wild-type Jurkat/sentinel/SKOV3-EGFRvIII). Each point represents aggregated data from all analyzed sections of one tumor. Horizontal bars indicate median values. Statistics: one-tailed paired t-test (**Methods**); P = 0.032 for the left two groups; P = 0.46 for the right two groups. N = 5 mice (left two groups), and N = 4 mice (right two groups). Connected dots denote contralateral tumors from the same mouse.

Circuit activation depended on the primary antigen (EGFRvIII) and all engineered circuit components. The full circuit, including EGFRvIII+ tumors, sentinels and auxin-gated effectors, showed an elevated CD69+ fraction in effector cells (**Fig. 6B**). By contrast, omission of any component of the circuit yielded similar levels of baseline activation. Within the same mouse (connected dots), the CD69+ fraction among CD3+ cells significantly increased in circuit-positive EGFRvIII+ tumors compared to contralateral tumors with wild-type THP-1 instead of sentinels (paired t-test, one-tailed, N = 5, p = 0.032) (**Fig. 6B**), consistent with spatially restricted activation of T cells to tumor sites. EGFRvIII-tumors or implants with wild-type THP-1 or Jurkat controls maintained similarly low CD69+ fraction, consistent with dependence on the engineered gates, and suggesting sentinel activation and auxin signaling remained localized (**Fig. 6B**). Taken together, these data indicate that engineered sentinel cells can locally license activation of Jurkat effector cells with HER2 CAR in vivo.

## DISCUSSION

Paracrine signaling is a central organizing principle in multicellular biology, enabling tissues to spatially gate potent cellular activities. Here, we established a fully synthetic paracrine (“synthacrine”) signaling system that operates in living tissue. Its features include a diffusible extracellular signal orthogonal to endogenous pathways, a sender density-dependent activation coverage, the ability to turn the signal on in response to defined inputs, and the ability to connect the received signal to defined outputs. Taken together, these features provide bio-orthogonality, tunability, and input-output modularity, and thereby enable synthetic, localized multicellular coordination in vivo.

The synthacrine system generates a spectrum of spatially organized responses. At low sender densities, we observed thin halos of localized activity around small sender cell clusters. As sender density increases, this transitions to contiguous millimeter-scale regions representing the summation of signaling fields of many individual sender clusters (**Fig. 2C, 2F–G**). Interestingly, a similar spatial overlap-driven transition has been proposed in tumor immunity, where widespread IFNγ signaling emerges only once dense T cell infiltration causes localized cytokine niches to overlap^26^.

The synthacrine system functions locally, even in the complex in vivo environment. Auxin signals generated within large, millimeter-scale regions did not measurably affect receivers in distant regions of the body, and systemic auxin remained at or below background-detection limits (**Fig. 2B, Fig. S1A**). This indicates that one can tune the strength and contiguity of a response within a tissue by modulating the density of auxin-producing cells (**Fig. 2E–G)** without activating responses in distal tissues.

The auxin signaling length scale in vivo was shorter than the centimeter-range gradients demonstrated in vitro^13^. This difference likely reflects features of the tissue microenvironment: e.g., transport barriers, cellular uptake, extracellular binding/partitioning, and especially clearance through vascular/lymphatic exchange. Together, these factors could yield an efficient auxin “sink” even without an engineered degradation module. Future work across additional tissues and disease states should clarify how this effective range scales with vascularization, extracellular matrix composition, cell packing, and immune infiltration, and whether engineered sinks or transport modules could further sharpen or extend signaling neighborhoods as needed.

As cell therapies grow in complexity, synthacrine channels could provide coordination across cell types and enable multicellular functions not possible with a single engineered cell type alone, including formation of functional niches or multicellular systems that collectively detect, amplify, and respond to complex input signals in a spatially distributed manner. Coupling auxin production to EGFRvIII-sensing in THP-1 cells allowed engineering of sentinel cells that regulate CAR in engineered Jurkat cells in a spatially restricted manner (**Figs. 3-6**). Engineerable human-orthogonal proteases such as TEVp allow flexible coupling and inversion of auxin signaling to indirectly activate CAR activity. This design required large, matching dynamic response ranges for the auxin-to-protease and protease-to-CAR interactions, which we achieved through iterative engineering (**Figs. S3B, D-G**). Further, it ensured tight suppression of CAR outside auxin-labeled regions (**Fig. 6B**).

In engineering the system, we addressed several challenges. First, auxin production initially occurred at lower rates in THP-1 cells compared to CHO-K1, possibly reflecting more limited expression capacity or tryptophan precursor abundance (**Fig. S2**). We addressed this through alternative designs of the auxin biosynthetic cassette (**Fig. S2**). Second, as mentioned, we circumvented the limitation of negative regulation by auxin-inducible degrons, by incorporating an intermediate protease step (**Fig. 4**). This strategy enabled tunable potentiation of CAR signaling by auxin. However, future designs that enable positive protein-level regulation by auxin could be more direct, compact, and modular. Third, a major challenge was achieving strong regulation of CAR activity by auxin. To enable this, we incorporated leucine zippers to colocalize the CAR and TEVp, along with multiple TEVp cleavage sites across several designs, and identified one configuration with strong cleavage efficiency but reduced auxin responsiveness (**Fig. S3D-F**). Further, we adopted 3 tandem AID repeats on protease, which helped to achieve a strong dynamic range for auxin-regulated CAR. However, additional AID repeats did not further increase dynamic range, suggesting the benefits of this mechanism have been optimized (**Fig. S3F, G**). Finally, once the system was engineered, we identified expression regimes that maximized the overall dynamic range of the sentinel-effector system, which occurred with lower levels of TEVp expression (**Fig. S3B**). These observations point to the importance of precise protein expression, which could be achieved through recent synthetic miRNA control circuits^27,28^.

Looking ahead, the designs introduced here could be generalized to provide new capabilities. Incorporating auxin transporters such as AUX1 and PIN2 for import and export, respectively, could improve signaling efficiency and tighten gradients. Since exogenous IAM provides a useful control knob to modulate auxin production rate (**Fig. S2**), future implementations could leverage IAM to tune the spatial range of auxin signaling in vivo. Engineering auxin-dependent auxin production could propagate auxin signals over longer ranges analogous to mechanisms used by the Nodal morphogen^29^. While we focused on restricting T cell activation, similar approaches could be used to spatially restrict anti-inflammatory cytokines for autoimmune disease, or modulate the inflammatory properties of tumor microenvironments. Similarly, instead of detecting tumor-specific antigens, sentinel cells could be reprogrammed to recognize markers of tissue/disease contexts by altering the antigen-sensing SNIPR domain. While we focused on sender cells (sentinels), which can actively regulate auxin production in response to local tissue state, it is also possible to imagine producing auxin within hydrogels to mark tumor margins, using viral vectors with iaaM and iaaH to label brain regions, or using ultrasound to spatiotemporally control auxin production^30,31^. Finally, additional circuits that allow sentinel cells to homeostatically control their own population densities and spatial arrangements could help to achieve predictable, scale-specific paracrine control in complex in vivo environments. These circuits could also incorporate mechanisms for active removal of auxin, analogous to the way natural cytokine systems such as IL-2 regulate spatial range through cell-based ligand removal^32^.

## METHODS

### Gene constructs

Constructs were assembled via Gibson cloning^33^ (New England BioLabs, M5520AAVIAL), restriction enzyme-based cloning, or were directly synthesized from GenScript. DNA fragments for cloning were prepared from existing plasmids using PrimeSTAR Max DNA Polymerase (Takara Bio, R045A), synthesized as gBlock (Integrated DNA Technologies), or through restriction enzyme digestion. Fragments were derived either from existing plasmids that have already been shown to be expressed in mammalian cells, or codon-optimized using the Integrated DNA Technologies codon-optimization tool. Some plasmid components, including backbones, were amplified from vYM155 (pSLCAR-CD19-BBz), which was a gift from Scott McComb (Addgene plasmid # 135992)^34^. Leucine zippers and some TEV cleavage sites are amplified from CMV-TO-nTVMVPteZtecTVMVP, which was a gift from Michael Elowitz (Addgene plasmid # 116082)^25^. Some HER2 scFvs were derived from pHR_SFFv_4D5-High-CAR_RHL002, which was a gift from Wendell Lim (Addgene plasmid #164825)^35^. Exact sequences used can be found in **Supplementary Data**.

### Tissue culture

All cells were cultured in tissue culture-treated plates or flasks. CHO-K1 cells, THP-1 cells, SKOV3 cells, and Jurkat cells were from ATCC. HEK293-FT cells were from ThermoFisher. Media were filtered through 0.22 µm filters after adding all supplements. Media formulations for culturing cells were as follows:

- CHO-K1 cells (and derivatives): Alpha-MEM supplemented with 10% FBS (Avantor), 1X GlutaMax (Gibco), and 1X pen/strep (Gibco).
- HEK293-FT: Dulbecco’s Modified Eagle Medium (Gibco) with 10% FBS (Avantor), 1X GlutaMax (Gibco), 1X pen/strep (Gibco), 1 mM sodium pyruvate (Gibco), and 1X non-essential amino acid (Gibco).
- SKOV3 cells (and derivatives): McCoy’s 5A modified with L-Glutamine (Gibco), with 10% FBS (Avantor), and 1X pen/strep (Gibco).
- Jurkat cells (and derivatives): Roswell Park Memorial Institute (RPMI) 1640 Medium with GlutaMax supplement and HEPES (Gibco), with 10% FBS (Avantor), 1X pen/strep (Gibco), and 1 mM sodium pyruvate (Gibco). Jurkat cells were maintained below 2 million cells/mL and split to 200,000 cells/mL when confluent.
- THP-1 cells (and derivatives): RPMI 1640 with HEPES and GlutaMax (Gibco), supplemented with 10% FBS (Avantor), and 1X pen/strep (Gibco). THP-1 cells were maintained between 200,000 cells/mL and 1 million cells/mL.

For co-culture experiments, media formats are experiment-specific (see relevant sections below).

### Lentivirus production and transduction

#### Lentivirus production

The ViraPower system (Invitrogen) was used to generate lentivirus, with modifications to the protocol. HEK293-FT cells were cultured for at least 1 week after thawing before virus packaging (kept in geneticin at 500 µg/mL). Cells were not passed beyond passage 16 and were not overgrown. On the day of transfection, the cells were 80-95% confluent.

For a T25 flask, media were gently replaced with 2 mL fresh culture media without geneticin. 14 µL Lipofectamine 2000 (Invitrogen) was added to 600 µL Opti-MEM (Gibco) and incubated for 5 min. In a separate tube, 3.6 µg of Virapower packaging mix (pLP1, pLP2, pLP/VSVG, total concentration is 1 µg/µL) and 1.2 µg of target plasmid (>100 ng/µL) were mixed in 600 µL Opti-MEM. The DNA mix was added to the Lipofectamine mix. The tube was gently inverted to mix, and incubated for 20 min. The mixture was added dropwise to the media covering cells without disturbing the monolayer.

The media were changed 8-16 hours later. Virus-containing supernatants were collected 48 h after the media change. Supernatants were centrifuged at 300 x g for 15 min or 500 x g for 10 min to remove cell debris in pellets, then used fresh or aliquoted (1 mL) and stored at −80 °C. Freeze-thaw cycles were avoided.

For a T75 flask, all volumes and amounts were scaled 2.5x relative to T25, except that 36 µL of Lipofectamine 2000 was used. When virus concentration was needed, the Lenti-X Concentrator protocol (Takara, PT4421-2) was followed.

#### Jurkat transduction

Lentiviruses were prepared as described. 20,000-50,000 Jurkat cells were prepared in 400 µL media/well in a 24 well plate. 200 µL lentivirus-containing HEK supernatant was added (for co-transduction with two viruses, 200 µL each). Volumes and cells can be scaled up for 12 well plates. Only for **Fig. S3C**, 400 µL 10X concentrated lentivirus was used. Polybrene (Sigma) was added to a final concentration of 8 µg/mL. Plates were gently shaken to mix. Cells were resuspended in fresh media 24 h later, and cultured for at least 48 hours before use. For specific conditions, refer to the raw data folder “readme” files.

#### THP-1 cell transduction

Lentiviruses were prepared as described, in T25 flask formats. Viruses were concentrated 5x by the Lenti-X concentrator (Takara). Transduction was enhanced by adding Polybrene (Sigma) at 5 µg/mL.

### Cell line engineering

All cells shown in this study were stably integrated with transgenes. CHO-K1 cells were stably engineered using the piggyBac transposon system and lentivirus system. SKOV3, Jurkat, and THP-1 cells were stably engineered by lentivirus transduction. Exact cells used can be found in **Supplementary Data**.

CHO-K1 sender cells were reused from the previous paper, with key plasmids available on Addgene^13^. CHO-K1 receiver cells used in **Fig. 2B** were also taken from the previous paper^13^, with the additional integration of BFP markers by PiggyBac (vYM136). CHO-K1 receiver cells used in **Fig. 2C-E** and **Fig. 2G** were transduced with lentivirus (1:6 diluted supernatant) expressing BFP (vYM226) instead.

THP-1 sentinel cells were co-cultured with EGFRvIII+ SKOV3 cells (mScarlet+), then sorted for positive EGFP expression. Specifically, cells were co-cultured for 96-120 h, then gently pipetted to lift cells (mostly THP-1), harvested and sorted (mScarlet negative, EGFP positive). After sorting, cells were further cultured and passaged for more than a week for complete recovery in preparation for downstream assays. Constitutive THP-1 auxin sender cells were also sorted using Citrine (Ver1) or EGFP (Ver2) markers.

Jurkat cells used in **Figs. S3C-G** were analyzed without sorting (gated on CAR-positive cells based on corresponding fluorescent markers). TEVp is not gated to allow presentation of “CAR only” condition as positive control. For **Fig. S3D-F**, TEVp positivity (EGFP+) was typically ~70%, but may be underestimated, as we observed CAR suppression without GFP positivity. For **Fig. S3G**, EGFP+ frequency was ~ 40% (P2-18), and ~60% (P2-17), but suppression remained strong. If EGFP was gated to be positive, P2-17 and P2-18 are similar in dynamic range (data not shown). It was observed that CAR suppression activity can be detected in cells that are not EGFP+, when CAR and TEVp are co-transduced. Effector Jurkat cells are used after sorting (described in **Fig. S3B**). To generate the effector cells in **Fig. 4-6**, Jurkats are first co-transduced with lentiviruses containing CAR and/or TEVp, and then transduced with lentivirus containing BFP as a constitutive marker. Lastly, the cells are sorted twice (described in **Fig. S3B**). The Jurkat cells with CAR/TEVp/BFP with low TEVp expression were used in **Fig. 5-6**.

### Flow cytometry

Adherent cells were detached using trypsin (0.25% for SKOV3 and CHO-K1) and neutralized by adding culture media. Jurkat and THP-1 cells were resuspended by thorough pipetting. Cells were then resuspended in Hank’s Balanced Salt Solution (Gibco) supplemented with Bovine Serum Albumin (2.5 mg/mL), or Cell Staining Buffer (Biolegend). For staining, Cell Staining Buffer (Biolegend), or 0.2% BSA in DPBS (Gibco) was used. Cells were passed through 40 µm strainers (Falcon) before acquisition. Flow cytometry was performed on a MACSQuant VYB (Miltenyi) or CytoFLEX S (Beckman Coulter) flow cytometer. Data were processed using EasyFlowQ (EasyflowQ v1.6)^36^ or FlowJo v10/v11 (BD Biosciences). Thresholds for the CD69 marker were based on wild-type Jurkat control groups (usually kept at 5% after thresholding to account for baseline noise).

### Cell Sorting

For sorting cells with different TEV protease expression, Jurkat cells were resuspended in Easysep Buffer (STEMCELL 20144) between 1-5 million cells/mL. Bulk sorting was performed on SONY MA900 using 100 μm chips, with sample pressure around 4. Cells were collected into tubes containing Jurkat culture media with extra FBS (final media contains 20% FBS) during sorting, then transferred to normal culture media for downstream cell culture and recovery.

### Animals

NSG mice (The Jackson Laboratory) were used. The study cohort included mice aged 7 weeks to 11 months. We did not observe obvious age-dependent differences in the data. Mice were housed on a 12 h light/dark cycle, and animals have ad libitum water and access to food. All experiments were performed with a protocol approved by the Institutional Animal Care and Use Committee of California Institute of Technology.

### CHO-K1 sender/receiver xenograft for flow cytometry and LC-MS

For each tumor, a total of 5 million cells were injected subcutaneously in a 1:1 mix of PBS and Matrigel (Corning, 356231/356230). Tumors were harvested 12-13 days after injection. Mice were euthanized and blood was collected by cardiac puncture (serum retained for LC-MS, related to **Fig. S1**). Tumors were dissected for homogenization and flow cytometry (related to **Fig. 2**).

### Tumor homogenization and flow cytometry

For flow cytometry, each tumor was minced with scissors and incubated in 10 mL digestion media (DMEM containing 2 mg/mL collagenase P and 0.1 mg/mL DNAseI) in a 15 mL conical tube at 37 °C for 2 h with continuous rotation. A 1 mL aliquot was filtered through a 40 µm strainer. Cells were kept at 4 °C and analyzed on the MACSQuant VYB Flow cytometer, Miltenyi.

### Histology and confocal imaging of CHO-K1 cell xenograft, and image processing

#### Histology

The conditions to generate subcutaneous CHO-K1 xenografts are the same as those xenografts used for flow cytometry and LC-MS. For CHO-K1 xenograft imaging (**Fig. 2C-E, andy 2G**), NSG mice underwent cardiac perfusion with PBS followed by 10% neutral buffered formalin (NBF). Xenografts were dissected and fixed in 10% NBF for 24 h at 4 °C. If tumors were not processed immediately, samples were washed once with PBS and stored in PBS or PBS with 0.05% sodium azide. Fixed xenografts were washed 3 times with PBS and cryoprotected with 30% (w/v) sucrose in PBS at 4 °C for 24-48 h on a rotator (5 mL tubes, protected from light) until samples sank. Samples were embedded in OCT and frozen on dry ice with isopropanol or kept in a −80 °C freezer. Tissue blocks were equilibrated to −17 °C to −20 °C in the cryostat chamber and sliced into 80 µm free-floating sections into PBS-containing untreated tissue-culture dishes. Remaining tissue blocks were returned to −80 °C. Sections were washed 3x with PBS and mounted onto slides. Prolong Diamond Antifade Mounting Media (Invitrogen, P36970) were applied for mounting. Slides were stored in the dark overnight before imaging and stored at 4 °C short-term.

#### Confocal imaging and image processing

Imaging was performed on a Zeiss LSM980 confocal microscope (20x lens, Plan-Apochromat 20x/0.8 M27). Images were acquired as tile scans and stitched automatically in the imaging software (ZEN blue v3.2). The data were then analyzed with Imaris (v11, Oxford Instruments) (for machine-learning based segmentation of cells, for **Fig. 2D**), and/or in-house Python scripts (for **Fig. 2C-G**) (annotated analysis pipeline in **Supplementary Data**).

For all xenograft analyses used in Fig. **2C-G**, samples could come from different flanks of the same mouse (2 xenografts/mouse), while being treated as independent based on the lack of contralateral cross-talk (**Fig. 2B** and **Fig. S1A**).

For **Fig. 2C** and **2G**, receiver pixels were defined as BFP+ (Channel 3 > 5,378 a.u.; threshold calibrated using the 99.99th percentile of BFP intensity from a sender-only negative control region) and plotted as a “plasma” heat map of mCherry intensity (Channel 1; linear map, 0–12,700 a.u.; over-range values indicated at the top of colormap as white). Sender pixels were defined as GFP+/BFP-/mCherry-(Channel 2 > 9,330 a.u., Channel 3 ≤ 5,378 a.u., Channel 1 ≤ 2,000 a.u.; with the GFP threshold calibrated from a receiver-only (sender-negative) control region (99.9th percentile)). The sender pixel map was Gaussian-smoothed (σ = 2.5 μm) to yield a local sender density map (area fraction; grayscale 0-0.1; over-range values indicated as red). Dotted lines indicate sender-rich areas for visual comparison. The rightmost panel (3.6% sender) of **Fig. 2G** and **Fig. 2C** are different zoom-in regions of the same section image.

For **Fig. 2D**, we quantified sender density–receiver response relationship from images of xenograft sections (senders and receivers segmented by Imaris v11, Oxford Instruments), computed in 200 µm × 200 µm windows. Per window, sender density (#/µm^2^) and median receiver mCherry were calculated; windows with ≥50 receivers were retained. Window-level values were log-binned by sender density (12 bins; ≥10 windows/bin) and plotted at geometric bin centers on log–log axes, with one curve per sample instance (sender concentration from 0.36%-10%). Most curves represent independent samples (N = 7 total different samples); one sample is shown as two curves from anatomically distinct xenograft regions. No-sender controls (receiver-only, 0% sender, and 0.01% sender seeded with sender-free fields by visual inspection) are shown in black to denote baseline mCherry; rare detected “senders” likely reflect classifier noise near the detection limit, whereas all other curves are colored.

For **Fig. 2F** analysis, briefly, receiver regions were defined by thresholding the BFP channel (Channel 3 > 5,378 a.u.), using a fixed cutoff calibrated from receiver-negative tissue. To quantify auxin-dependent repression, we computed the fraction of receiver pixels with low mCherry (Channel 1 < red_threshold). Red_threshold was set using sender-free (receiver-only) control xenografts: for each control image, we extracted mCherry intensities from BFP+ receiver pixels and took the 5th percentile (to account for fluctuation of mCherry expression in CHO-K1 receiver cells in vivo, or non-receiver artifact pixels bright in BFP channel); red_threshold was then chosen as the minimum of these per-image percentiles (i.e. 2,083 a.u.) and applied unchanged to all samples to calculate the fraction of receiver pixels that are responsive (% Receiver area responsive) (annotated analysis pipeline in **Supplementary Data**).

### LC-MS analysis of auxin concentration in blood serum

#### Serum preparation

Whole blood (100–200 µL) was collected by terminal cardiac puncture. Blood was allowed to clot at room temperature, then centrifuged at >15,000 x g for 30 min at 4 °C (without pre-chilling the centrifuge). The clear upper serum layer (typically ~50% of the whole blood volume; may appear pink/red) was collected and stored at −20 °C.

#### LC-MS sample preparation

This protocol was modified from previous literature^37^. Thawed serum (50 µL) was mixed with 150 µL pre-chilled acetonitrile, vortexed for 4 min at room temperature, and incubated for 10 min at −20 °C to precipitate proteins. Samples were centrifuged at 16,128 ×g for 20 min at 4 °C. Clean supernatant (100 µL) was transferred to an autosampler vial for LC-MS.

#### Standard curve

IAA was spiked into FBS and serially diluted. For each standard, 50 µL of the standard was mixed with 150 µL acetonitrile, and serial dilution was performed by transferring 50 µL into 150 µL acetonitrile repeatedly. Series 1: 25,000 ng/mL, 6,250 ng/mL, 1,563 ng/mL. Series 2: 200 ng/mL, 50 ng/mL, 12.5 ng/mL, 3.1 ng/mL. Standards from both series and an FBS-only control were processed using the same LC-MS preparation steps (starting from vortexing 4 min at room temperature).

#### Positive controls

IAA was dissolved in serum from wild-type untreated mice at 0 ng/mL, 50 ng/mL, and 6,250 ng/mL, then processed as above (starting from vortexing for 4 min at room temperature).

#### LC-MS

LC-MS is set up based on literature^37^ and performed on ACQUITY I-Class UPLC System (Waters) comprised of a binary solvent manager (BSM), a flow-through needle autosampler (FTN), and thermostated column compartment connected with Xevo G2-S QTof mass spectrometer (Waters). MassLynx and Quanlynx (v4.1, Waters) were used to acquire and analyze data. ACQUITY UPLC HSS T3 1.8 µm (2 mm x 100 mm, Waters) column was used for chromatographic separation. Mobile phase A is water + 0.1% formic acid, and mobile phase B is acetonitrile. The flow rate was constant at 0.3 mL/min. The gradient program began at 2% B, rose to 60% B by 2.2 min, and was held there until 3.5 min. The program then returned to 2% B at 4 minutes and ended at 5 minutes. The injection volume was 2 µL. The ion source was operated in positive ion mode, capillary voltage 2.5 kV, with desolvation gas flow of 900 L/hr at 450 °C. The mass axis was calibrated using sodium formate and lock mass correction was performed using protonated leucine enkephalin at 556.2771 m/z.

### Construction of EGFRvIII-responsive SNIPR

We constructed the synthetic EGFRvIII sensing receptor following the modular design described in a previous publication^38^. Briefly, the scFv (clone 139) against EGFRvIII was connected to the optimized version of CD8α hinge, human Notch1 transmembrane domain, human Notch2 juxtamembrane domain (vYM201), and then Gal4-VP64 transcription activation domain to form a fusion protein. The sequence is codon-optimized, synthesized and cloned to a third-gen lenti-expressing plasmid. scFv sequence is provided in **Supplementary Data**.

### Optimization of iaaH, iaaM cassette

#### Co-culture of sentinel cells (THP-1) and target (SKOV3) cells

SKOV3 cells were seeded at 25% (for 5 day co-cultures) or 50% (for 2 or 3 day co-cultures) confluence, and allowed to settle in the incubator for around 4 hours. Culture media were then removed and THP-1 cells were seeded at 1:1 (THP-1 to SKOV3) ratio into the co-culture system. We used a 1:1 mix of THP-1 and SKOV3 media for any co-cultures. For imaging experiments, the seeding ratio is adjusted to 1:5 (THP-1 to SKOV3). Imaging was done using the same inverted Olympus IX81 microscope setup as previously described^13^, with the same Zero Drift Control, automated stage, camera (Andor iKon-M CCD, 13 µm/px), 20x objective, light source, and control software. 2×2 binning was used.

#### Co-culture of sentinel cells (THP-1) and reporter (CHO-K1) cells

THP-1 and mCherry-AID/mTagBFP expressing CHO-K1 cells were seeded in a 1:1 mix of THP-1 and CHO-K1 media at a ratio of 5:1. IAM was supplied as indicated. At the end of co-culture, both cells were collected and analyzed by flow cytometry with the addition of 2,000 CountBright beads (Life Technology C36950) for counting the cells.

#### Inferring auxin level in conditioned media from standard curve

Conditioned media were harvested, and centrifuged at 500 x g for 5 minutes to clear out any remaining cells. An equal amount of CHO-K1 fresh media was then mixed in before applying to CHO-K1 reporter cells. In parallel, CHO-K1 reporter cells were seeded in a 2x titration curve of IAA ranging from 0.01 µM to 25 µM or 100 µM, depending on the expected range of auxin, serving as a standard curve. The inferred concentration of IAA in the conditioned media was calculated by comparing to the standard curve and factoring in the 2x dilution.

### Engineering the auxin-responsive TEVp and the TEVp-responsive CAR

#### Jurkat T cell activation assay for evaluating CAR-TEVp designs

Co-culture cell conditions were as indicated in **Fig. 4** and **Fig. S3**. SKOV3 (or derivatives) were plated on day 0. On day 1, Jurkat cells (or derivatives) were co-cultured with SKOV3 cells (or derivatives) at an effector:target ratio of 1:2-1:5 (refer to raw data folders for exact ratio) in Jurkat media and incubated for 24 h. Jurkat cells were harvested on day 2 by gentle but thorough pipetting, followed by staining and flow cytometry to quantify CD69 expression.

For **Fig. S3C-G**, Jurkat cells in the auxin condition were pre-incubated with auxin on day 0 for 24 h before co-culture, and auxin treatment was continued during co-culture. For **Fig. S3A** and **Fig. 4**, Jurkat cells in the auxin condition were treated with auxin only during the 24 h co-culture starting on day 1.

### Staining for CD3 and CD69 in co-cultures

Jurkat and/or THP-1 cells were resuspended from SKOV3 co-cultures by gentle but thorough pipetting. Cells were washed by centrifugation (500 x g, 5 min) to remove media. After two washes in the staining buffer (PBS + 0.2% BSA, or Biolegend Cell Staining Buffer), cells were stained in 50 µL staining buffer with 2.5 µL CD69 antibody (Biolegend 310910) and/or CD3 antibody (Biolegend 300431), and incubated on ice for 15-20 min in the dark. For **Fig. 4** and **Fig. 5**, after incubation, 130 µL media were added and mixed before spinning down. For all experiments in **Fig. 4, Fig. 5** and **Fig. S3**, cells were washed twice after incubation. Then, cells were resuspended in 200 µL of the staining buffer. For **Fig. S3C-G**, the final staining buffer contained 1,000x diluted SYTOX Blue Dead Cell Stain (Invitrogen). For **Fig. 4, 5**, and **Fig. S3A**, all T cells were labeled with constitutive BFP, thus SYTOX Blue was excluded. Cells were passed through a 40 µm strainer and analyzed by flow cytometry. When SYTOX Blue was added, cells were incubated for 5 min before acquisition. If not immediately acquired, stained cells were stored in the dark at 4 °C for up to 8 h.

### In vitro full circuit assay

On day 0, 25,000 SKOV3 cells (or derivatives) and 25,000 THP-1 cells (or derivatives) were plated in one well of a 48-well plate in 300 µL Jurkat media, with designs specified in **Fig. 5**. On day 1, plates were spun (500 x g, 5 min). Media (255 µL) were removed and replaced with 255 µL Jurkat media containing 6,250 Jurkat cells (or derivatives) and IAM (final concentration 50 µM). On day 4, CD3 and CD69 staining was performed and Jurkat/THP-1 cells were analyzed by flow cytometry, after resuspension by gentle but thorough pipetting.

### Histology and confocal imaging of full circuit tumors and image processing

#### Tumor injection and treatment

On day 0, mice received subcutaneous injections of 500,000 THP-1 (or derivatives), 500,000 SKOV3 (or derivatives), and 500,000 Jurkat (or derivatives), as described in **Fig. 6**, in 1:1 PBS/Matrigel (Corning, 356231/356230). Mice received 0.5 mg/mL IAM via PBS for drinking instead of water from day 4 to day 7; the IAM solution was made fresh and replaced daily. Mouse appearance and weight were monitored to prevent potential dehydration and discomfort. Mice were euthanized on day 7 by CO_2_.

#### Tumor freezing and cryo-sectioning

Tumors were snap-frozen in OCT-filled cryomolds (**Fig. 6**) by placing the molds in the vapor phase of liquid nitrogen. 10 µm sections were cut from the cryostat and directly mounted onto slides, air-dried for 10-30 min at room temperature, and stored at −80 °C.

#### Immunostaining and imaging

Sections were fixed in 10% NBF for 15 min at room temperature, washed 3x with PBS to remove NBF and an additional 3x with PBS to fully remove OCT, and blocked in 5% BSA in PBS for 1 h at room temperature on a shaker, protected from light. CD3 antibody (Biolegend 300431) and CD69 antibody (Biolegend 310910) were each diluted 1:5 in 5% BSA in PBS and applied to sections. Samples were incubated for 1 h at room temperature in a humidified chamber, washed 3x with PBS, mounted with Prolong Antifade Diamond Mounting Media (Invitrogen, P36970), and imaged on a Leica Stellaris confocal microscope (20x objective, HC PL APO CS2 20x/0.75 IMM). The images were acquired as tile scans and automatically stitched in the imaging software (LAS X). Imaging analysis was performed using Imaris (Oxford Instruments) for cell segmentation/classification, and in-house Python scripts for quantification. The threshold for CD69+ was set as mean infrared intensity per segmented object (> 5,000 a.u.) to yield near-zero CD69+ percentage in negative control group (wild-type Jurkat/sentinel/SKOV3-EGFRvIII sample) (**Supplementary Data**). For statistical analysis, we performed Shapiro-Wilk tests on the within-mouse paired differences and found no evidence to reject the null hypothesis that these differences were normally distributed (**Supplementary Data**). Further, one-tailed paired t tests were used for prespecified directional comparisons: the localized-circuit model predicts greater CD69 activation in the functional-circuit flank than in the matched non-functional flank within the same mouse, and in vitro data (**Fig. 5** and **Fig. S4**) predicted higher background CD69 activity in the control circuit with effector cells than the control circuit with wild-type Jurkat cells. Representative images were processed using FIJI (CD3/blue channel minimum-maximum: 0-1,000 a.u., CD69/infrared channel minimum-maximum: 0-15,823 a.u., determined based on control group stained without primary antibodies) (**Supplementary Data**).

### Statistics

Measurements of data points are taken from distinct samples, unless otherwise noted.

### Curve fitting with Hill function

Data in **Fig. 2E, S1A, S2A** are fitted using a repressive Hill function equation with background. They have different x and y definitions, but the form remains consistent:

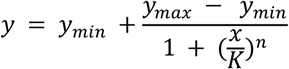

For **Fig. 2E**,

*x* = *Fraction of sender seeded in the xenograft*

*y* = *Median mCherry intensity*(*a. u*.) *for receiver pixels within a xenograft section*

*y*_*min*_ = *Inferred mCherry intensity* (*a. u*.) *with saturating presence of senders*

*y*_*max*_ = *Inferred mCherry intensity* (*a. u*.) *with no senders*

*n* = *Hill coefficient*

*K* = *EC*_50_

For **Fig. S1A**,

*x* = *Auxin concentration* (µ*M*)

*y* = *Median receiver mCherry intensity* (*a. u*.)

*y*_*min*_ = *Inferred mCherry intensity* (*a. u*.) *with saturating presence of auxin*

*y*_*max*_ = *Inferred mCherry intensity* (*a. u*.) *with no auxin*

*n* = *Hill coefficient*

*K* = *EC*_50_ (µ*M*)

For **Fig. S2A**,

*x* = *Sender number*

*y* = *Median receiver mCherry intensity* (*a. u*.)

*y*_*min*_ = *Inferred mCherry intensity* (*a. u*.) *with saturating presence of senders*

*y*_*max*_ = *Inferred mCherry intensity* (*a. u*.) *with no senders*

*n* = *Hill coefficient*

*K* = *EC*_50_

Data in **Fig. 3C** and **4D** are fitted using an activating Hill function equation, with background. They have different x and y definitions, but the form remains consistent:

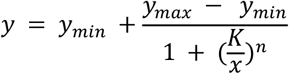

For **Fig. 3C**,

*x* = *Fraction of EGFRvIII positive SKOV*3 *cells*

*y* = *Inferred auxin concentration in media* (*μM*)

*y*_*min*_ = *Inferred baseline auxin concentration in media* (*μM*) *with no EGFRvIII positive SKOV*3

*y*_*max*_ = *Inferred auxin concentration in media* (*μM*) *with only EGFRvIII positive SKOV*3

*n* = *Hill coefficient*

*K* = *EC*_50_

For **Fig. 4D** (each effector group),

*x* = *Auxin concentration* (*μM*)

*y* = *Activated T cell* % (*CD*69 *positive*)

*y*_*min*_ = *Inferred activated T cell* % (*CD*69 *positive*) *with no auxin*

*y*_*max*_ = *Inferred activated T cell* % (*CD*69 *positive*) *with saturating concentration of auxin*

*n* = *Hill coefficient*

*K* = *EC*_50_ (*μM*)

For exact parameter values and parameter fitting strategy, see individual processing pipeline for details (**Supplementary Data**).

## Use of generative AI in code drafting

Portions of the code used for data quantification, analysis, and plotting (analysis scripts and FIJI macros) were drafted with assistance from ChatGPT (OpenAI; GPT-4/GPT-5). All code was subsequently reviewed, edited, and validated by the authors. The authors take full responsibility for the content of this publication.

## DATA AND CODE AVAILABILITY

- All original and processed data have been deposited at CaltechDATA as **Supplementary Data** (data.caltech.edu) and are publicly available at the time of publication (DOI: 10.22002/e79gg-zch92).
- The code used for analyzing the data is included in the same folder in **Supplementary Data** (data.caltech.edu) and will be publicly available at the time of publication (DOI: 10.22002/e79gg-zch92).
- Any additional information required to reanalyze the data reported in this paper is available from the lead contact upon request.

## MATERIALS AVAILABILITY

Plasmids generated in this study will be deposited to Addgene. All unique/stable reagents generated in this study are available from the lead contact upon completion of a materials transfer agreement. For further information and requests for resources and reagents, please contact the lead contact, Michael B. Elowitz.

## ACKNOWLEDGEMENTS

We thank Jordi Garcia-Ojalvo, Ellen Rothenberg, Sarkis Mazmanian, Matt Thomson, Neda Bagheri, Allison Li, Rong Lu, Wang Yuan-Hsi, Jonathan Hoang, Dan Piraner, Yanruide Li, Richman Lee, Sicheng Li, Lena Gamboa, John Ngo, and Andrew McMahon for insightful scientific input and technical suggestions; Rongrong Du, Jee Won Yang, Shirin Shivaei, James Linton, Leah Santat, Yuxing Yao, Zhiyang Jin, Hao Shen, Evan Mun, Sheng Wang, Lucy Chong, George Daghlian, Justin Lee, Tom Duan, Ishaan Dev, Mohamed Abedi, Bo Gu, Dongyang Li, Shiyu Xia, Yodai Takei, Martin Tran, Ron Hadas, Victoria Tobin, Andrew Lu, Felix Horns, Michael Flynn, Duncan Chadley, Dhiraj Indana, Gal Manella, Jacob Parres-Gold, Ben Emert, Julio Revilla, Di Wu, Rui Malinowski, Jo Leonardo, and all other members of the Elowitz lab and Shapiro lab for critical feedback and administrative support; Scott McComb, Crystal Mackall, Louai Labanieh, Wilson Wong, and Sarah Adams for sharing experimental materials; imaging was performed in the Biological Imaging Facility, with the support of the Caltech Beckman Institute and the Arnold and Mabel Beckman Foundation; and Inna-Marie Strazhnik for graphical design. LC-MS analyses were performed in the Resnick Water and Environment Laboratory at the California Institute of Technology. This publication is based on research supported by the National Institutes of Health (*EB030015*) and the G. Harold and Leila Y. Mathers Charitable Foundation (*12540447*). M.B.E. and M.G.S. are both Howard Hughes Medical Institute Investigators. The funders had no role in study design, data collection and analysis, decision to publish, or preparation of the manuscript. This article is subject to HHMI’s Open Access to Publications policy. HHMI lab heads have previously granted a nonexclusive CC BY 4.0 license to the public and a sublicensable license to HHMI in their research articles. Pursuant to those licenses, the author-accepted manuscript of this article can be made freely available under a CC BY 4.0 license immediately upon publication.

## AUTHOR CONTRIBUTIONS

K.L., Y.M., M.G.S., and M.B.E. conceived and designed the study. M.G.S. and M.B.E. supervised the study. H.R.L., H.L., M.L., M.B.S., N.F.D., A.S.F., E.C-H., and A.L. performed or assisted with experiments and data analysis. K.L. and M.B.E. wrote the manuscript with input from all authors.

## DECLARATION OF GENERATIVE AI AND AI-ASSISTED TECHNOLOGIES IN THE WRITING PROCESS

During the preparation of this work, the authors used ChatGPT (OpenAI; GPT-4/GPT-5) in order to improve the readability and language of the manuscript. After using this service, the authors reviewed and edited the content as needed. The authors take full responsibility for the content of the published article.

## Supplementary Information

**Figure S1** relates to the CHO sender/receiver module (**Fig. 2**).

**Figure S2** describes THP-1 constitutive sender optimization, supporting **Figure 3.**

**Figure S3** is related to **Figure 4**.

**Figure S4** is relevant to **Figure 5**.

**Figure S1:**
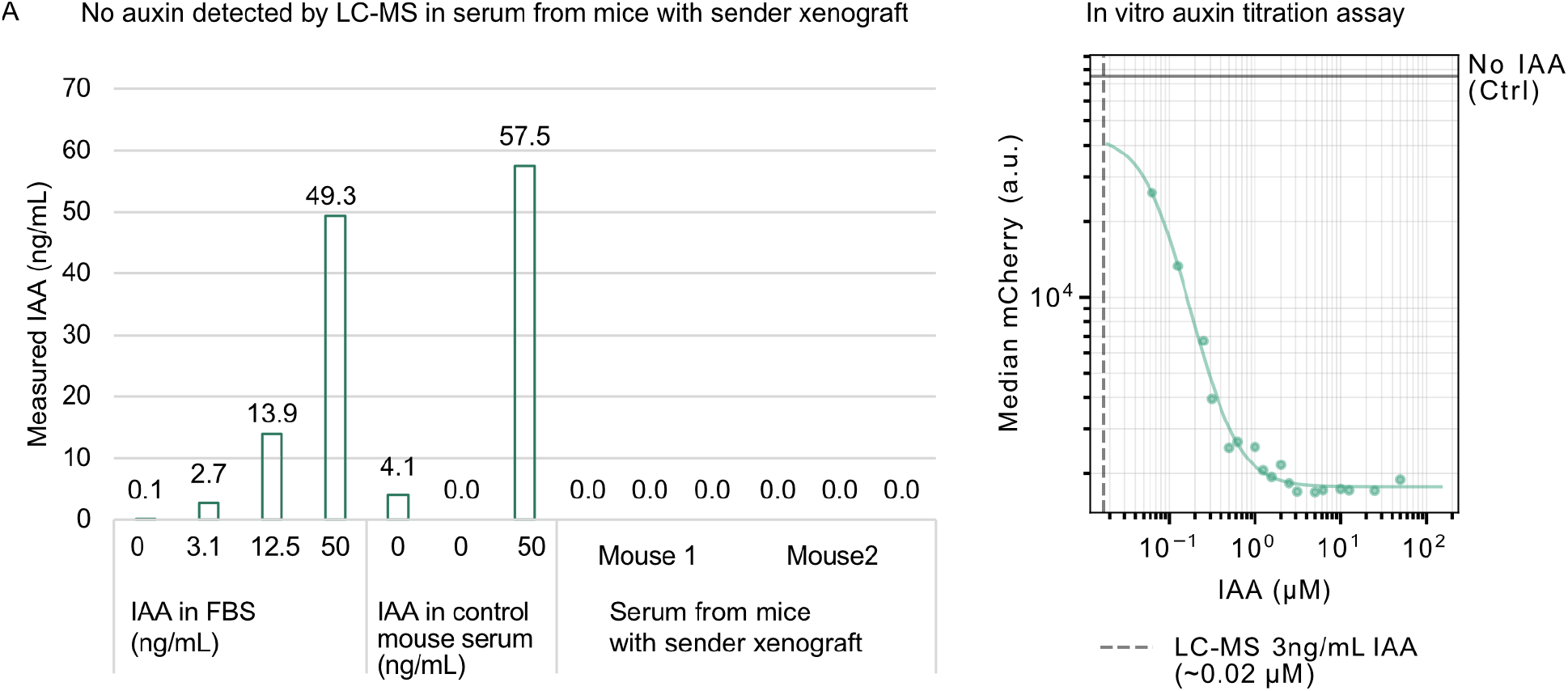
Auxin remains largely undetectable in serum. (**A**) (Left) Serum auxin was measured by liquid chromatography-mass spectrometry (LC-MS) from mice bearing xenografts containing 10% senders/90% receivers and was negligible. As controls, FBS and serum from receiver-only mice were spiked with defined IAA concentrations. Controls were N = 1 per matrix; CHO sender Mouse1 and Mouse2 were biological replicates, each measured in technical triplicates. An IAA spike of 3 ng/mL corresponds to ~0.02 µM, and is insufficient to elicit a strong receiver response (right). Note that in raw data, all samples had some peak area signal, but some were shown as 0 because the area values were at/below the background level, as determined by the calibration curve (**Supplementary Data**). Hill function was used to fit data points (**Methods**). See **Supplementary Data** for fitted parameters.

**Figure S2:**
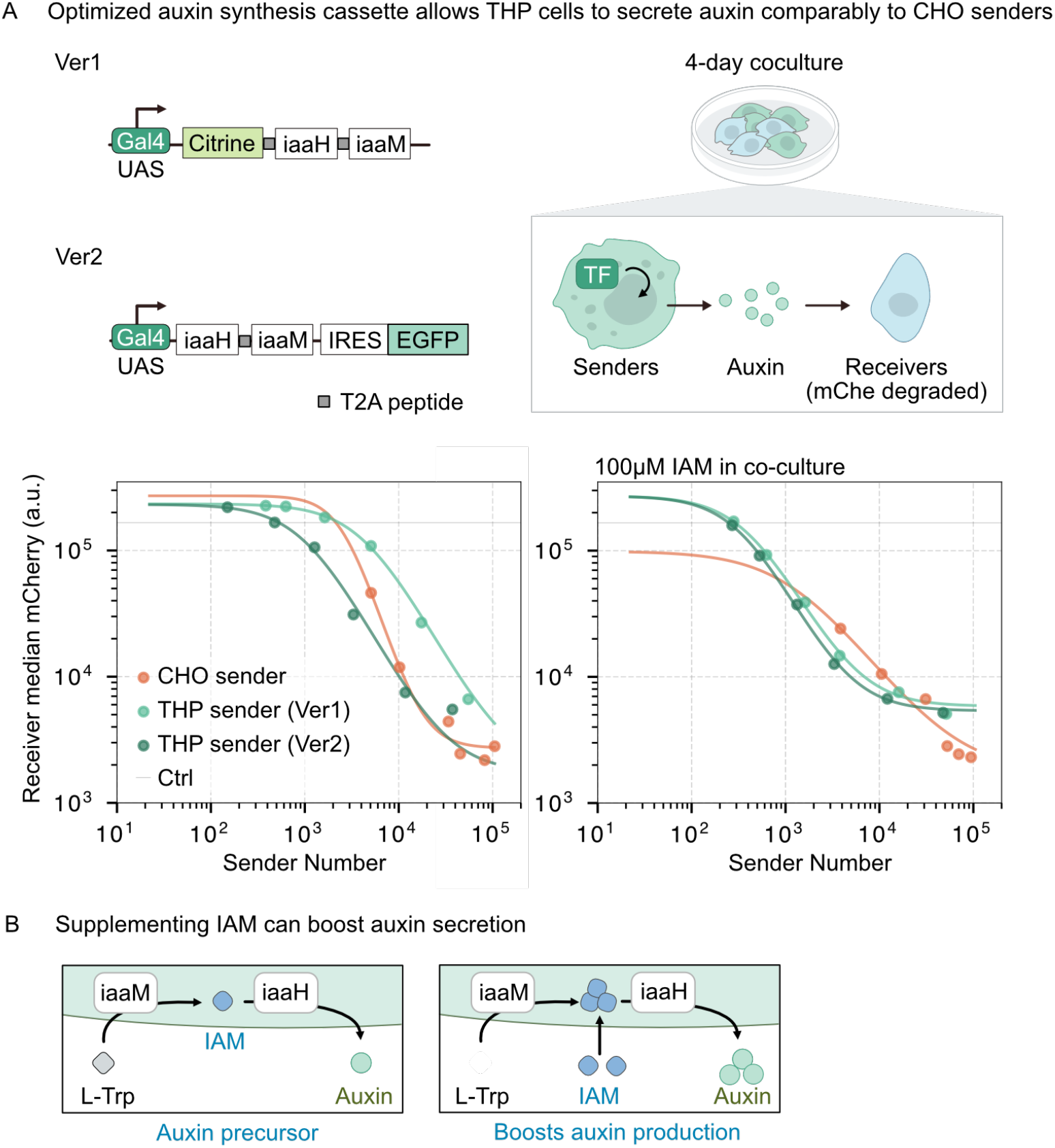
Optimized auxin production construct allows constitutive THP auxin senders to secrete auxin at levels comparable to CHO senders. (**A**) Different constitutive auxin senders (Ver1/Ver2) were co-cultured with CHO auxin receivers for 4 days. The cells then underwent flow cytometry to compare auxin secretion strength (receiver mCherry-AID degradation). IAM was applied in co-culture to investigate if the improved performance in the optimized construct is due to improved iaaM activity. Hill function was used to fit data points (**Methods**). See **Supplementary Data** for fitted parameters. N = 1 for each concentration group for the titration curves. (**B**) IAM, as the auxin precursor, is supplied to boost auxin production.

**Figure S3:**
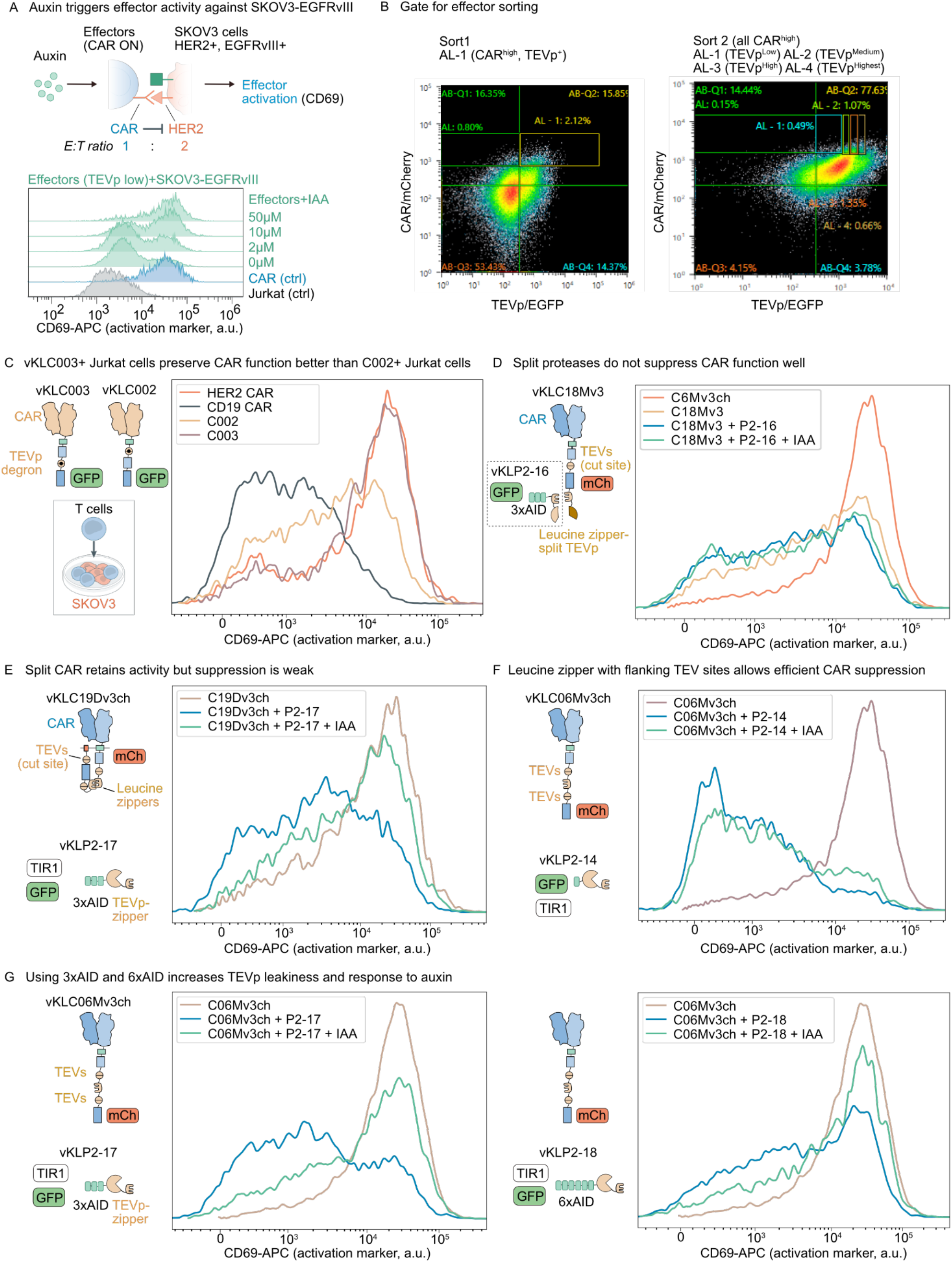
Effector design iteration, additional validation and expression optimization by sorting. (**A**) Same co-culture setup as in **Fig. 4D**, but using EGFRvIII+ SKOV3 cells to examine auxin-gated CAR activity in the presence of EGFRvIII. (**B**) Gate used for effector sorting. (**C-G**) Different CAR/TEVp designs leading towards the final design. Jurkat cells were transduced with corresponding lentiviruses encoding the constructs, co-cultured with SKOV3 cells in a 1:2-1:5 ratio. The cells were pre-treated with auxin for 24 h before another 24 h co-culture with SKOV3 cells. The cells were gated on CAR+ based on the corresponding construct-linked fluorescence marker (EGFP or mCherry). IAA, indole-3-acetic acid (auxin). N = 1 for each example.

**Figure S4:**
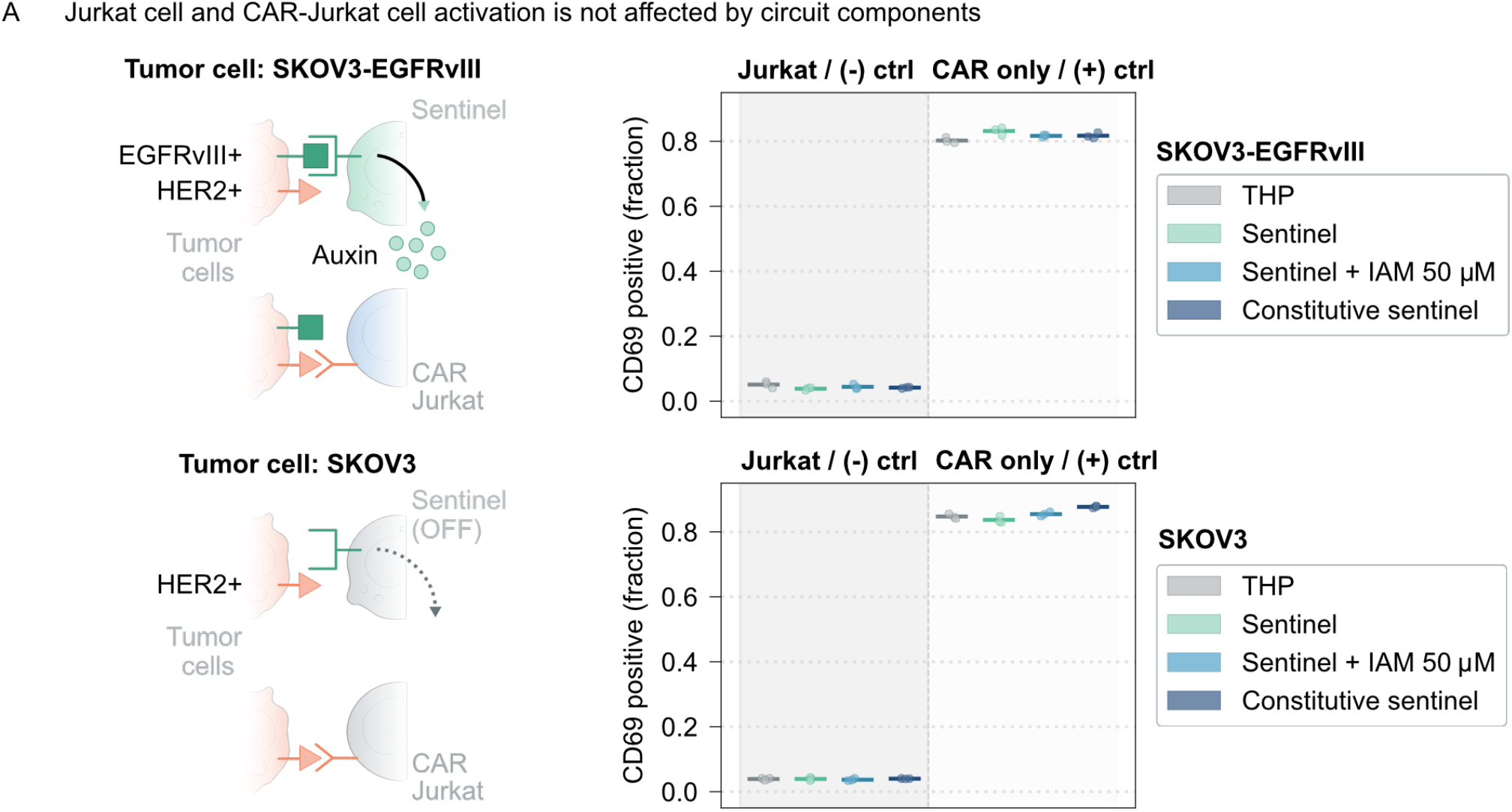
The activation of CAR-only Jurkat and wild-type Jurkat cells is independent of the presence of EGFRvIII or auxin production. (**A**) The co-culture and readout setup was the same as **Fig. 5A**, but with Jurkat effector cells replaced by CAR Jurkat cells without TEVp or wild-type Jurkat cells, as control groups. N = 3 for each condition.

